# The bacterial chaperone CsgC inhibits functional amyloid CsgA formation by promoting the intrinsically disordered pre-nuclear state

**DOI:** 10.1101/2025.03.21.644623

**Authors:** Anthony Balistreri, Divya Kolli, Sanduni Wasana Jayaweera, Daniel Lundahl, Yilin Han, Lily Kalcec, Emily Goetzler, Rachel Alessio, Brandon Ruotolo, Anders Olofsson, Matthew R. Chapman

## Abstract

*E. coli* assembles a functional amyloid called curli during biofilm formation. The major curlin subunit is the CsgA protein, which adopts a beta-sheet rich fold upon fibrillization. The chaperone-like protein CsgC inhibits CsgA amyloid formation. CsgA undergoes a 3-stage aggregation process: an initial lag phase where beta-rich nuclei form, an exponential elongation phase, and a plateau phase. It is currently not known if CsgC inhibits amyloid formation by inhibiting formation of a pre-fibril nucleus, or if CsgC inhibits a later stage of amyloid formation by blocking monomer addition. Here, CsgC homologs from *C. youngae*, *C. davisae*, and *H. alvei* were purified and characterized for their ability to interrogate CsgA amyloid formation. Each of the CsgC homologs prolonged the lag phase of *E. coli* CsgA amyloid formation similar to *E. coli* CsgC. Additionally, we found *E. coli* CsgC interacted transiently and weakly with a monomeric, pre-nucleus species of CsgA which delayed amyloid formation. A transient CsgC-CsgA heterodimer was observed using ion mobility-mass spectrometry. When CsgC was added to actively polymerizing CsgA, exponential growth commonly associated with nucleation-dependent amyloid formation was lost. Adding preformed CsgA seeds did not rescue exponential growth, indicating that CsgC also has inhibitory activity during fibril elongation. Indeed, CsgC interacted strongly with CsgA fibers, suggesting the interaction between CsgC and CsgA fibers can slow new fiber growth. CsgC displays unique inhibitory activity at multiple stages of amyloid formation. CsgC acts as an energy-independent chaperone that transiently interacts with prefibrillar CsgA and an amyloid fiber.

## Introduction

Amyloids are fibril protein aggregates that are characterized by their distinct morphology, stability, and tinctorial properties (1). Abnormal accumulation of amyloids can lead to a wide range of pathologies, including diseases such as Parkinson’s Disease, Alzheimer’s Disease, and Type II Diabetes (2). The proteins that comprise amyloid fibers are often intrinsically disordered, or have intrinsically disordered regions, which adopt a repetitive cross-β strand structure upon fibrilization (2). Amyloid intermediates are cytotoxic and have been shown to be more toxic than mature amyloid fibers (3, 4). While inhibiting disease-associated amyloid formation remains a significant focus of research, a growing body of work is focused on utilizing functional amyloids as a model for amyloid formation and regulation.

Functional amyloids are a class of amyloid that provide a useful function for the cell, the list of which continues to grow (5–7). The most widely studied functional amyloid is curli, the main protein component of the *E. coli* biofilm matrix (6). Curli fibrils surround cells in the biofilm and provide protection from desiccation, predation, and viral infection (6). The *E. coli csg* operons contain seven curli specific genes that function as the producer and regulator for curli formation, with each protein playing a significant role (8). The main amyloid protein, CsgA, is translocated into the periplasm by the SecYEG pore (6). Periplasmic CsgA is then shepherded to the nonameric CsgG outer membrane pore by CsgE, a periplasmic chaperone (9). Two auxiliary proteins, CsgB and CsgF, are also exported through the CsgG pore and act as a nucleator and anchor, respectively, to allow CsgA amyloid fibers to form and attach to the cell’s outer surface (10, 11). Lastly, the operon contains a second periplasmic protein, CsgC, which functions as a potent inhibitor of CsgA and CsgB amyloid formation (12).

CsgC is a 110 amino acid protein with an immunoglobulin-like β-sandwich structure and proposed chaperone-like activity (13). *E. coli* CsgC is an efficient sub-stoichiometric inhibitor of amyloid formation against its native client *E. coli* CsgA (12). CsgC can also inhibit amyloid formation by *E. coli* CsgA homologs like the *Pseudomonas* amyloid FapC, and the human disease-associated amyloid alpha-synuclein (12). Little is known about the CsgC-client protein interaction, except i) the interaction is guided by electrostatic interactions (14) and ii) there is a weak client sequence determinant (12). Many details about CsgC-mediated amyloid inhibition remain poorly understood. CsgC is a simple protein with no predicted active sites, domains, or significant amino acids. CsgC requires no hydrolysable substrate, cofactor, or cellular energy to perform its activity. CsgC does not take up a substrate or produce a quantifiable product; the only evidence of CsgC activity is the inhibition of amyloid formation. Therefore, CsgC represents a potentially novel class of anti-amyloid chaperone proteins that do not utilize an energy source and do not maintain a stable interaction with the client protein.

Amyloid formation occurs in a series of steps, during which soluble, disordered monomers join an insoluble, ordered amyloid fiber. In a solution containing amyloid-competent monomeric proteins, a low abundance species arises from the pool of disordered monomers which acts as a nucleation site for fiber formation (15). This nucleation event represents the rate-limiting step for amyloid formation. Disordered monomers add on to these “nuclei” and begin the progressive and thermodynamically favorable process of forming amyloid fibers (15). Fiber formation marks the transition of amyloid proteins from the soluble state to the insoluble, and in the case of curli fibrils, a transition from an intrinsically disordered monomeric conformation to a stable and ordered β-sheet rich aggregate.

Previous work suggested that CsgC inhibits CsgA amyloid formation via electrostatic interactions (14), however, it is unclear what stage(s) of amyloid formation is inhibited by CsgC and what conformer of CsgA that CsgC binds to (14, 16–18). Here, we used several biochemical and biophysical techniques to provide direct evidence that CsgC can inhibit CsgA amyloid formation through two different mechanisms. We tested CsgC homologs from four Gammaproteobacteria and found that they inhibit *E. coli* CsgA amyloid formation kinetics through more than one mechanism. CsgC transiently binds a sub-population of monomeric CsgA on-pathway to form amyloid and prolongs the formation of the aggregation-prone β-sheet rich fold state characteristic of amyloid proteins. CsgC strongly binds CsgA fibrils and blocks fibril elongation. We propose that these chaperone-like mechanisms are in accordance with the *in vivo* function of CsgC: preventing formation of cytotoxic amyloid aggregates in the bacterial periplasm and acting as an amyloid fibril formation inhibitor.

### Experimental procedures

#### Bacterial Growth

All overnight cultures were grown in LB supplemented with 100 μg/mL ampicillin and/or 50 μg/mL kanamycin at 37 °C with shaking at 220 rpm. When necessary, LB plates were supplemented with ampicillin 100 μg/mL or kanamycin 50 μg/mL.

#### Strains and Plasmids

The full list of strains, plasmids, and primers can be found in the Supporting Information. Gibson Assembly was used to construct plasmids using NEBuilder® HiFi DNA Assembly Cloning Kit (Cat. No. E5520S). Primers were designed using the NEBuilder webtool (https://nebuilder.neb.com) and purchased from IDT (https://www.idtdna.com). Mutagenized plasmids were constructed in the MC1061 cell background. Correct mutations were confirmed using Sanger sequencing provided by Eurofins (https://www.eurofins.com/genomic-services/our-services/custom-dna-sequencing/). Plasmids were extracted from transformants using Promega PureYield™ Plasmid Miniprep System (Cat No. PRA1223). Miniprepped plasmids were transformed into an expression strain (NEB3016) for purification.

#### Protein Purification

CsgA was purified as described previously (19). CsgC and its variants were purified as described previously (20). Size exclusion chromatography was performed for CsgC purification as previously described (21). Briefly, Ni-NTA affinity chromatography elution fractions were pooled, concentrated, and passed through a 0.22 μm filter. The sample underwent gel filtration using a Superdex 75 10/300 GL column (Cat No.45-002-903) attached to an Äkta pure protein purification system. Elution fractions were assayed for protein concentration using A220 and/or Pierce™ BCA Protein Assay Kit (Cat No. 23225). Samples corresponding to elution peaks were analyzed using SDS-PAGE.

#### Denaturing Gel Electrophoresis and Dot Blot

Purified protein samples were diluted in 4X SDS loading buffer and run on a 15% SDS PAGE gel with 10 μL loaded into each well (21). The gels were stained with Coomassie blue dye to visualize protein bands. Dot blots were performed by spotting a nitrocellulose membrane with 2 μL of protein and allowing the spots to fully dry. Blots were blocked for one hour in a mixture of TBST and skim milk. Blocked blots were then washed three times with TBST and probed with a primary antibody solution against CsgA (1:12,000) (21) or CsgC (1:4000) (17).

Primary antibody was removed from blots which were then washed three times with TBST prior to the addition of a secondary antibody solution. Secondary antibodies against rabbit IgG and conjugated with IRDye 800CW (Cat No. NC9401842) were used to image the blots in a Licor Odyssey FC.

#### Surface Plasmon Resonance

Surface Plasmon Resonance (SPR) experiments were performed on a BIAcore™ 3000 (GE Healthcare) using a CM5 chip for the immobilization of sonicated CsgA fibrils and gel filtered CsgA monomers of a stable CsgA mutant (22). Immobilization was carried out following protocols from previous fibrillar SPR studies (23), with CsgA fibrils immobilized at 1300 RU and monomers at 500 RU. After immobilization, the flow cells were probed with gel-filtered CsgC at concentrations ranging from 4 μM to 0.25 μM (two-fold dilutions). CsgC was injected for 5 minutes over the CsgA fibrils and 7.5 minutes over the CsgA monomers. The experiment was conducted in phosphate buffer (pH 7.5) containing 0.05% Tween, at a temperature of 25°C. Prior to immobilization, CsgA fibrils were sonicated for 1 minute in a water bath.

#### Preparation of modified NHS columns

CsgC was purified as described previously (20). NHS-activated agarose beads were buffer exchanged into 50 mM potassium phosphate buffer (pH = 7.4) and reacted with purified CsgC (1mg CsgC/ml NHS-activated agarose beads, Cat. No. 26200) overnight at 4 °C with gentle rocking. The reacted resin was then washed with 50 mM potassium phosphate buffer, pH = 7.4 to remove any unbound CsgC and cleaved NHS-groups. Reaction progress was tracked by 260 nm and 220/280 nm absorbance values to measure increase in cleaved NHS-groups and decrease in free CsgC protein, respectively. SDS-PAGE and Pierce™ BCA Protein Assay Kit (Cat No. 23225) were used to ensure complete binding of CsgC to the resin. Unreacted NHS groups still bound to agarose beads were cleaved by addition of a quenching buffer (1M Tris HCl, pH 7.5) and incubated for 30 mins at room temperature with gentle rocking. The fully reacted resin was buffer exchanged into 50 mM potassium phosphate buffer, pH = 7.4 to remove quenching buffer. 260 nm and 220/280 nm absorbance values were measured to ensure complete quenching of unreacted NHS-linked agarose beads. Modified NHS columns were stored at 4 °C until use. Individual columns were only used once for each assay.

#### Sample Preparation for Ion mobility-mass spectrometry (IM-MS)

Purified CsgA and CsgC were buffer exchanged into 20 mM ammonium acetate (pH 7.4) using Thermo Scientific Zeba™ Spin Desalting Columns 7k MWCO (Cat No. 89892). The protein concentration after buffer exchange was assayed using Thermo Scientific Pierce™ Rapid Gold BCA Protein Assay Kit. CsgA and CsgC were each diluted to 20 µM with 20 mM ammonium acetate, mixed at 1:1 ratio. The mixture was incubated at 37°C for 23 hours. Time points were taken at 0 hr, 3 hr, 6hr and 23hr.

#### IM-MS

IM-MS data was collected on a quadrupole ion-mobility time-of-flight (TOF) mass spectrometer (Synapt G2 HDMS, Waters, Milford, MA, USA) with a nano-electrospray ionization (nESI) source. The source was operated at positive mode with the nESI voltage set at 1.0-1.2 kV, the sampling cone was set to 15 V and the bias was set to 42 V. The source temperature was set to 20°C. The traveling–wave ion mobility separator operated at a pressure of approximately 3.4 mbar with wave height and wave velocity set at 30V and 500 m/s, respectively. The m/z window was set from 100 – 8000 m/z with a TOF pressure of 1.5e-6 mbar. Mass spectra were analyzed using MassLynx 4.1 and Driftscope 2.0 software (Waters, Milford, MA, USA). CCS (Ω) measurements were externally calibrated using a database of known values in helium. We reported the standard deviations from replicate measurements of CCS and an additional ±3% to incorporate the errors involved in the calibration process.

#### Thioflavin T Binding Assay

Assays were performed as previously described (19). Briefly, freshly purified CsgA was diluted with phosphate buffer to 20 μM or 10 μM and combined with an excess of the amyloid-specific dye thioflavin-T (ThT) (Cat No. AC211760250). Amyloid formation was monitored by measuring an increase in ThT fluorescence at 495 nm (450 nm excitation). Assays were performed in triplicate at microscale within 96-well plates and measured with Infinite Pro M200 or Infinite Nano^+^ F200 Tecan plate readers. CsgC proteins were purified, diluted with phosphate buffer, and added to specified assays in the reported stoichiometric ratio. Fiber stimulating “seeds” were produced by obtaining previously purified WT CsgA fibers and were sonicated directly before adding to a specified assay. Time to half-maximum (T_1/2_) is calculated by the time at which the Thioflavin-T fluorescence reaches half of the maximum Thioflavin-T fluorescence. Duration of lag phase (T_lag_) was calculated as described by Arosio et al (24). Briefly, T_lag_ is calculated by T_1/2_ subtracted by half of the maximum of the first derivative of the sigmoidal ThT fluorescence.

#### Circular Dichroism

CsgA was diluted with phosphate buffer to a final concentration of 15 μM. Secondary structure was measured at room temperature every 60 minutes (in triplicate) with the JASCO-1500 using default parameters. Measurements were averaged, the baseline signal of phosphate buffer was subtracted, and the curve was smoothed using a Savitzky-Golay filter. Final curves were normalized by concentration and path length.

## Results

### E. coli CsgC and three CsgC homologs affect multiple stages in the CsgA amyloid formation Pathway

CsgC was first identified in *E. coli* (EC), but *csg* operons from other Gammaproteobacteria contain CsgC homologs (25). *Citrobacter youngae* (CY), *Cedecea davisae* (CD), and *Hafnia alvei* (HA) are three Gammaproteobacteria that have *csg* operons containing homologs to EC CsgC. These four CsgC homologs have similar structures, as predicted by Alphafold 2.0 (26), however, they share a range of sequence identity with CsgC EC: CY (69%), CD (51%), and HA (35%) (**Fig. 1A-D and Fig. S1)**. CsgC and each homolog was tested for inhibitory activity against *E. coli* CsgA using ThT binding assays. CsgC EC was the most efficient inhibitor of *E. coli* CsgA amyloid formation, with full inhibition at a 1:500 stoichiometric ratio (40 nM) (**Fig. 1E**). CsgC CY, CsgC CD, and CsgC HA all inhibited CsgA aggregation to a lesser extent (**Fig. 1F-H**). All ThT assay results were fit with a sigmoidal function curve. We derived the duration of the lag phase for each curve as a proxy for the nucleation rate of CsgA polymerization (24). The lag phase of *E. coli* CsgA was significantly prolonged upon addition of CsgC homologs in 11 of 12 assays (**Fig. 1I**). The lag phase was not determined in two cases because the ThT signal never increased within the time frame of the experiment (**Fig. 1I**). The only condition where addition of CsgC did not affect the lag phase was with addition of the lowest ratio (1:1000) of CsgC HA homolog, which has the lowest sequence identity to CsgC EC (**Fig. 1H** and **I**).

**Fig. 1.**
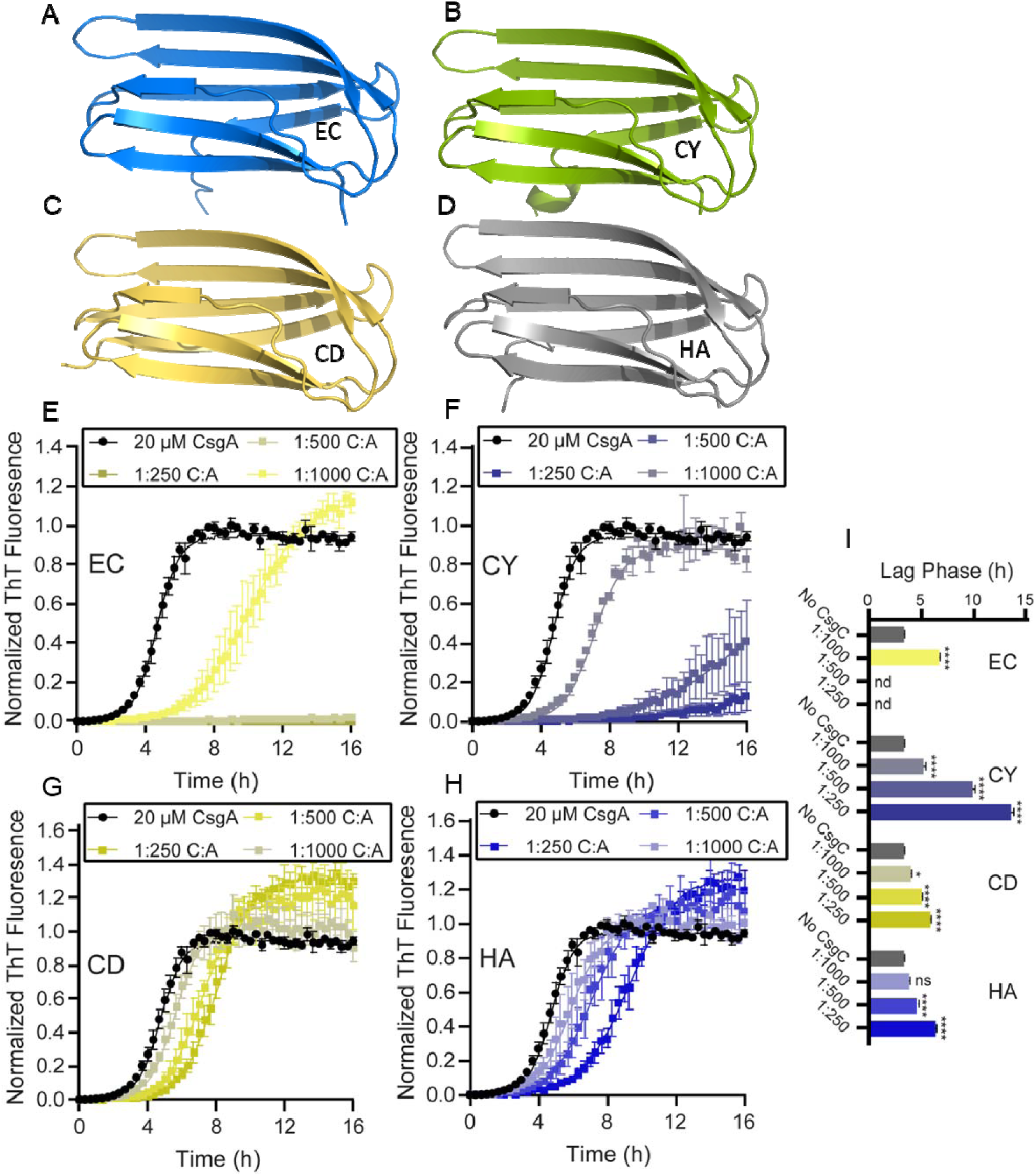
CsgC EC and its homologs are effective at extending the lag phase CsgA amyloid formation. **A-D)** AlphaFold 2.0 predicted structures for CsgC EC, CY, CD, and HA. **E-H)** 4 different CsgC homologs were purified and tested for their ability to inhibit CsgA in vitro. In all cases CsgA was freshly purified and mixed in the stated stoichiometric ratio with CsgC homologs. The data points shown represent the average of triplicate experiments, the error bars show the SEM, and the curves were fit using the sigmoidal logistic function described in (24). **I)** The calculated lag phase for each homolog according to the fit curves. The error bars represent the SEM for each value. “nd” was used to denote values which were not defined by the equation used. Significance was attributed using a one-way ANOVA analysis comparing all values to the uninhibited CsgA condition; *p < 0.05, ****p < 0.00005

Amylofit was used for additional analysis of the ThT curves produced in **Fig. 1**. Amylofit is a global fitting software that fits ThT binding assay curves, calculates estimated values for reaction orders, rate constants, and more to describe amyloid aggregation kinetics (27). Meisl and coworkers suggest that Amylofit can be used to determine the molecular species or stage along an aggregation pathway that is affected by an inhibitor (27). The rate expressions used to fit ThT curves describe the formation of CsgA amyloid fibers starting with unfolded CsgA monomers:

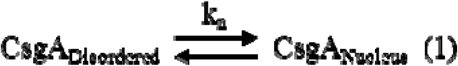

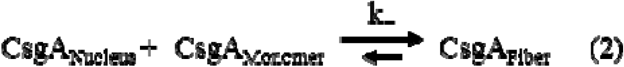

A primary nucleus forms in solution from an unknown number of folded CsgA monomers (k_n_, eq.1). After a critical concentration of nuclei are present, CsgA monomers use nuclei as folding templates, sparking rapid fiber elongation (k_+_, eq.2). For some amyloid proteins, the presence of amyloid fibers in solution stimulates new fiber growth (this could be described by rate constant k_2_).

We inputted the same data displayed in **Fig. 1E-H** into Amylofit and normalized each curve to reflect the final aggregate amount in the uninhibited CsgA condition. The “secondary nucleation dominated” model was used to fit the ThT curves because, under these aggregation conditions, this model provided the best fit. At first, all rate constant parameters (k_n_, k_2_, and k_+_) were allowed to be fit individually, providing the best fit curves for each condition (**Fig. S2A, S3A, S4A, and S5A**). Next, all parameters except one were set to a global constant using the values derived from CsgA fibril formation in the absence of inhibitor. This was repeated for all rate constant parameters. Using this method, each rate constant parameter was set to individually fit each inhibited condition (**Fig. S2B-D, S3B-D, S4B-D, and S5B-D**). This method illustrated the degree to which each rate constant can explain changes in the ThT curve upon addition of increasing concentrations of inhibitor (27). Best fit curves were created for each condition and the mean squared residual error (MRE) results from each condition provided a global metric for goodness of fit for all curves (27). Low MRE values are an indication that the fitted rate constant is affected by the addition of the inhibitor. Interestingly, the rate constant that provided the lowest MRE value was different between each homolog, but MRE values were generally close to the universal best fit MRE value (**Fig. S2E, S3AE, S4E, and S5E**). The rate of secondary nucleation (k_2_) and elongation rate (k_+_) often resulted in lower MRE values than the primary nucleation rate (k_n_), suggesting that CsgC could be affecting CsgA aggregation at the level of fibril elongation. Taken together, **Fig. 1-2** and **Fig. S2-5** indicated that CsgC and its homologs may affect CsgA amyloid formation at multiple stages.

**Fig. 2.**
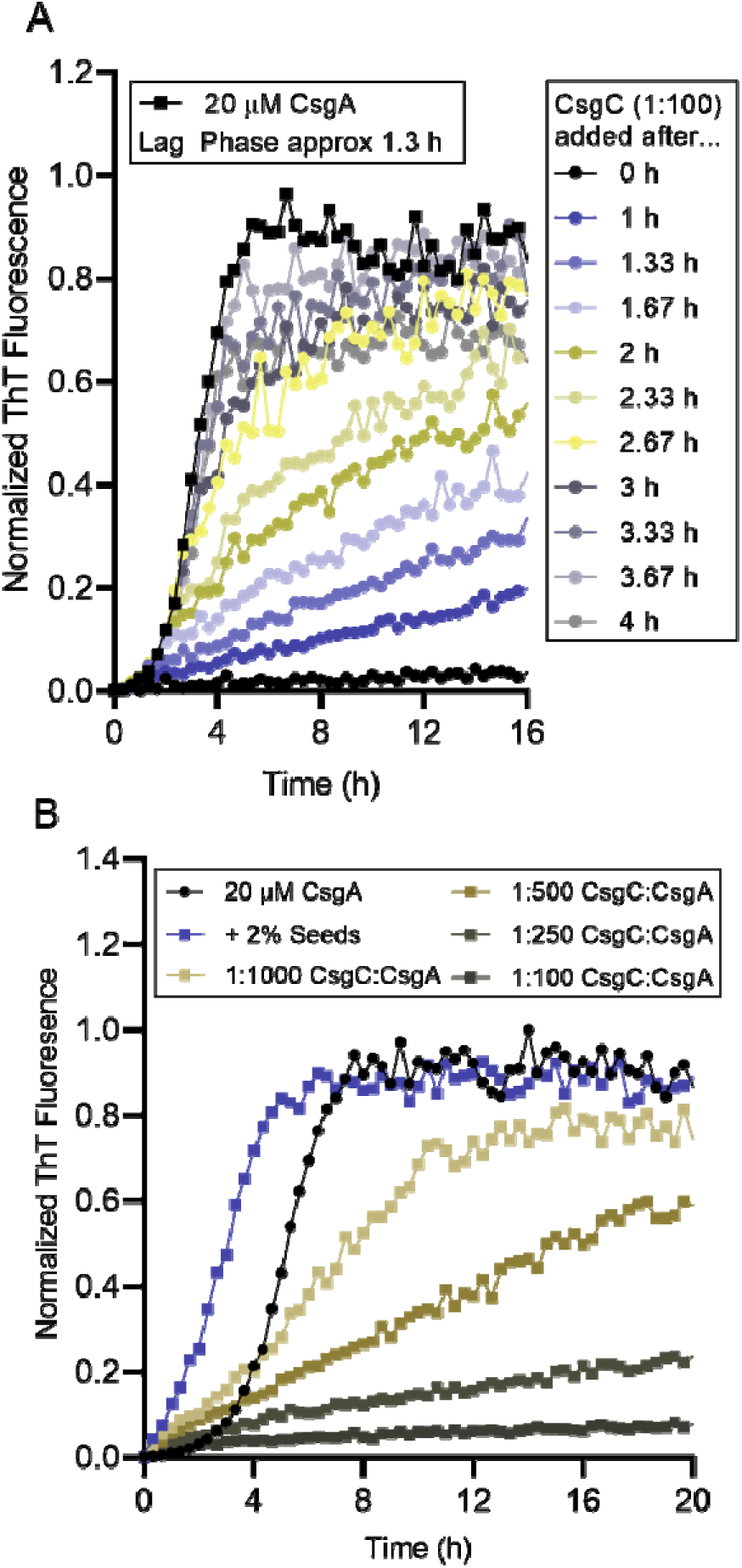
CsgC inhibits amyloid formation during the fiber elongation stage. **A)** 200 nM (1:100) CsgC was added to actively polymerizing CsgA at the times indicated and amyloid aggregation was monitored in a ThT binding assay. **B)** In a similar assay, CsgC was added to a 2% seeded reaction of CsgA at various ratios as indicated in the key above (All olive conditions also contain 2% seeds).

### CsgC partially inhibits CsgA amyloid formation during the elongation phase

CsgA amyloid formation follows a primary nucleation-dependent polymerization mechanism that has three distinct phases: an initial lag phase, an elongation phase, and a stationary phase. The elongation phase is marked by fibril elongation directed by soluble protein monomer addition onto fiber ends (28, 29). Importantly, even during the elongation phase there are intrinsically disordered CsgA monomers still in solution capable of either primary nucleation or addition onto a fiber (30).

CsgC was added at different time points to actively polymerizing CsgA to determine how CsgC inhibitory activity changes throughout amyloid formation. First, CsgC was added to a final concentration of 200 nM (1:100 CsgC:CsgA molar ratio) at different time points after the start of CsgA amyloid formation (0 hours) up to 4 hours (**Fig. 2A**). When CsgC was added at time 0 hours the ThT signal remained very low compared to CsgA samples without CsgC, indicating nearly total inhibition. When CsgC was added to CsgA after 1 hour, the ThT signal increased linearly (**Fig. 2A**). When CsgC was added after the lag phase had ended, the result was a linear increase in ThT signal followed by a gradual decrease reaching a plateau in signal (**Fig. 2A**) indicating that CsgC was capable of slowing fiber elongation. However, it is not clear whether the inhibition is due to prevention of new nuclei formation or prevention of fibril elongation.

To determine if CsgC inhibitory activity is solely based on preventing nuclei formation, CsgC was added to CsgA that had been seeded by the addition of preformed CsgA fibers (**Fig. 2B**). During a seeded reaction, CsgA is presented with preformed fibers and the fibers act as a template for amyloid formation, effectively diminishing the need for primary nucleation and the lag phase entirely (**Fig. 2B, dark blue squares compared to black circles**) (31). When CsgC was added to a seeded CsgA reaction, the result was a linear increase in ThT signal (**Fig. 2B**). In both cases (**Fig. 2A** and **B**), the addition of CsgC into solution when fibril formation is largely dependent on CsgA monomer addition to actively growing amyloid fibrils resulted in partial inhibition of amyloid formation indicating that CsgC inhibited fibril elongation.

### CsgC binds to CsgA fibrils and not to CsgA monomers

Surface plasmon resonance was used to determine the binding kinetics between CsgC and CsgA monomers or CsgA fibrils. Surface plasmon resonance can interrogate interactions between proteins and calculate a K_D_ (32). Either CsgA fibrils (**Fig. 3A**) or CsgA monomers (**Fig. 3B**) were bound to a CM5 chip using standard amino-coupling reaction. A double cysteine variant of CsgA that retains an intrinsically disordered structure under oxidizing conditions was used to prevent bound monomeric CsgA from aggregating during the course of the SPR experiment (22). Solutions containing freshly purified monomeric CsgC were flowed over the sensors.

**Fig. 3.**
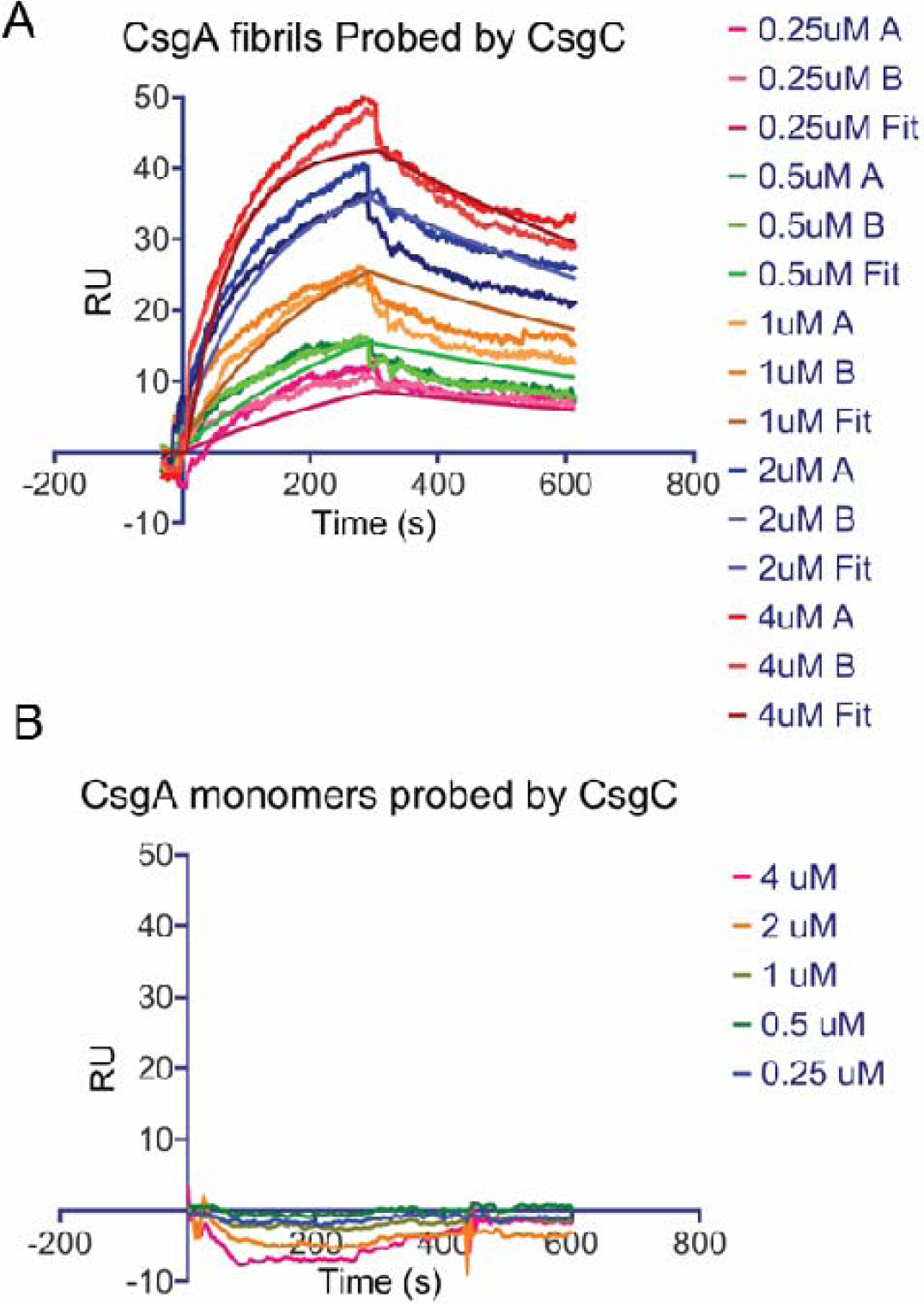
Sensograms of CsgA fibrils and monomers probed by CsgC. **A)** Interaction between CsgA fibrils and CsgC, showing an equilibrium dissociation constant (K_D_) of 360 ± 10 nM. The concentrations labeled “A” represents the first injection, while “B” denotes the secondary injection. The fitted curve represents the curve fitting performed using Scrubber2. **B)** No detectable interaction between CsgA monomers and CsgC across all tested concentrations. Residual standard deviation = 3,194 RU.

The sensogram data with curve fits is shown in **Fig. 3**. The residuals indicated a generally good fit, with only minor deviations observed at certain concentrations. Notably, concentrations below 250 nM were excluded from the kinetic analysis due to detection limitations, as the responses at these lower concentrations were too weak to be reliably measured by SPR. Sensogram data indicated that monomeric CsgA had no measurable binding to monomeric CsgC (**Fig. 3B**). However, monomeric CsgC interacted with CsgA fibrils with a dissociation rate of K_d_ = (1,18 ± 0.05) × 10 ³ s^-1^ and an equilibrium dissociation constant of K_D_ = 360 ± 10 nM (**Fig. 3A**).

### Transient interaction between monomeric CsgC and non-fibrillar CsgA is sufficient to delay amyloid formation

Previous studies have shown that CsgA incubated with CsgC does not readily form higher order fibrillar aggregates (14, 17). This data, in conjunction with data in **Fig. 1** and **Fig. S2-5**, indicated that CsgC can also inhibit the initial conversion of monomeric CsgA into an oligomeric amyloid. Due to the undetectable binding between monomeric CsgC and CsgA from **Fig. 3**, we utilized a modified pull-down assay to determine if CsgC interacts transiently with prefibrillar CsgA.

*N*-<u>h</u>ydroxy-<u>s</u>uccinimide (NHS) resin beads were coupled to CsgC protein for the modified pull-down assay. Another batch of *N*-<u>h</u>ydroxy-<u>s</u>uccinimide (NHS) resin beads were quenched with Tris to use as a control. Immediately after Ni-NTA and SEC purification, a CsgA sample was split into equal parts. One aliquot of the purified CsgA was left on ice, while the other aliquot was passed through a column containing either the CsgC-linked beads or the Tris quenched beads (**Fig. 4A**). CsgA eluent from the column was then diluted into the wells of a ThT binding assay to monitor amyloid formation. The CsgA from the Tris-quenched resin condition began aggregating at 2 hours, consistent with CsgA that had been left on ice (**Fig. S6A**). Interestingly, the CsgA that passed by the CsgC-linked column had a diminished propensity to form amyloid, displaying an approximately 77 min increase in ThT lag phase and a 75 minute increase in the time it takes to reach half maximum ThT fluorescence compared to CsgA that had been left on ice (**Fig. S6B**). CsgA eluted from the CsgC-linked column displayed a 51 minute increase in the lag time when directly compared to CsgA eluted from the Tris-quenched column and a 52 min increase in the time it takes to reach half maximum ThT fluorescence, displaying that transient interaction between CsgA and CsgC is sufficient for delayed amyloid formation **(Fig. 4B** and **D)**. Buffer was eluted from the Tris-quenched column and the CsgC-linked column and added into a ThT binding assay with freshly purified CsgA to ensure that no CsgC, NHS, or Tris eluted from the resin that could affect CsgA aggregation (**Fig. 4C**).

**Fig. 4.**
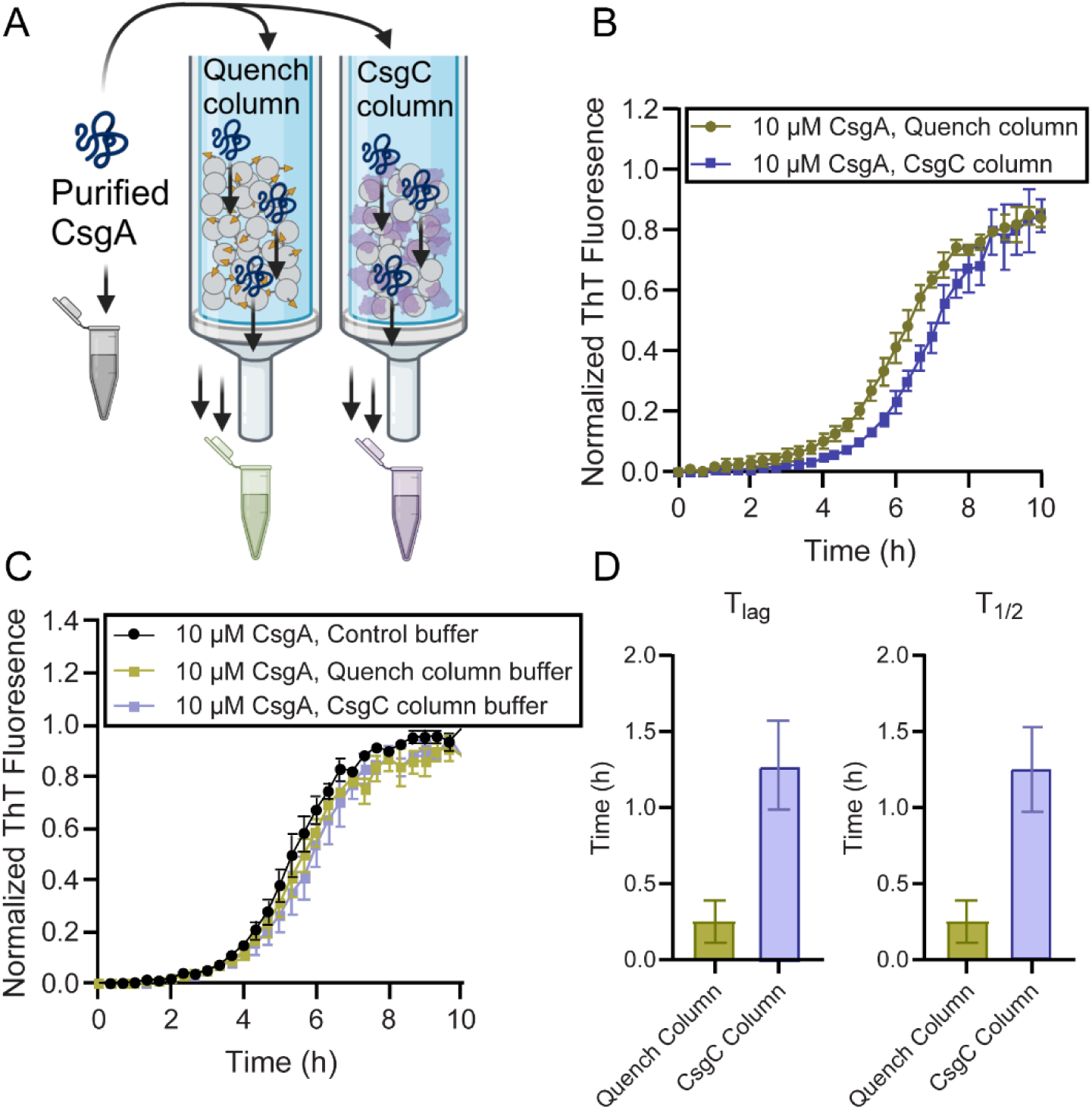
Passing monomeric CsgA by immobilized CsgC delayed the formation of ThT positive aggregates. **A)** Affinity purified CsgA was passed through a column of NHS agarose resin that was either amino coupled to CsgC or Tris base to quench the free NHS binding spots. **B)** Eluent CsgA was then diluted to 10 µM in phosphate buffer for a ThT binding assay to observe amyloid aggregation kinetics. **C)** Quenched and CsgC resin was incubated in phosphate buffer for 24 hours. The incubated phosphate buffers were used to dilute freshly purified CsgA to 10 µM and added to a ThT binding assay. **D)** A measurement of the lag phase (T_lag_) and time to half maximum fluorescence (T_1/2_) was calculated using a sigmoidal curve function. Error bars represent the standard error of the mean across three technical replicates.

CsgA protein concentration was quantified prior to, and following, elution from each resin to determine how much CsgA was retained on the CsgC-linked beads. Little to no difference was observed between the eluent concentration of CsgA eluted from the control Tris-linked resin beads (31.2 μM) and the eluent concentration of CsgA eluted from the CsgC-linked resin beads (25.7 μM). To determine if the small difference in concentration was due to CsgA retention on the CsgC-linked resin column, the CsgC-linked resin and Tris-linked resin was treated with hexafluoroisopropanol (HFIP) to digest CsgA aggregates left on the resin beads. Samples were then analyzed by a dot blot and SDS-PAGE to determine if any CsgA is retained on the beads following elution. Both the SDS-PAGE gel and the dot blot indicated that no detectable CsgA was retained on the resin following elution (**Fig. S7**). This further supports that CsgC transiently interacts with CsgA, and that this interaction is sufficient to observe the extended lag phase in **Fig. 4**.

Additionally, the change in CsgA secondary structure from disordered to beta-sheet rich was visualized using circular dichroism **(Fig. S8)**. Amyloid protein folding is often assessed using CD by measuring the formation of a spectral minimum at 220 nm indicating the formation of a beta-sheet rich and amyloidogenic secondary structure. No significant effect of CsgC transient interaction on beta-sheet structure formation was detected by circular dichroism. With the indication of an interaction between CsgC and CsgA monomers by ThT but not by CD, we decided to look more closely at their interaction using a highly sensitive native mass spectrometry technique.

### A transient 1:1 CsgC:CsgA complex is detected

Native mass spectrometry (native MS) uses electrospray ionization conditions to preserve non-covalent protein-protein and protein-ligand complexes (33). The added dimension of the ion mobility (IM) spectrometry allows ions to be separated based on their size, shape, and charge (**Fig. 5A**) (34). We purified both CsgC monomers and CsgA monomers, added them together in a 1:1 mixture, and analyzed the resulting solution for any interactions between the two proteins. During our first time point, we observed monomeric and dimeric species of both proteins, including low abundance heterodimeric complexes of CsgA and CsgC (**Fig. 5B** and **C**). CsgA exhibited a broad charge state distribution ranging from 5+ to 13+, as typically observed for intrinsically disordered proteins (**Fig. 5B**) (35). The CsgA:CsgC mixture remained soluble throughout the 23 hour time course while apo CsgA, serving as a control sample, formed micro-scale aggregates that led to the clogging of the nESI emitters and prevented the collection of both 6 hour and 23 hour time point data. An analysis of collisional cross-section (CCS) values of CsgA monomers showed very little difference between time 0 and 3 hour (**Fig. S9A**). Similarly, we also saw no changes in CCS for CsgC monomer ions through the 23 hour time course (**Fig. S9B**).

**Fig. 5.**
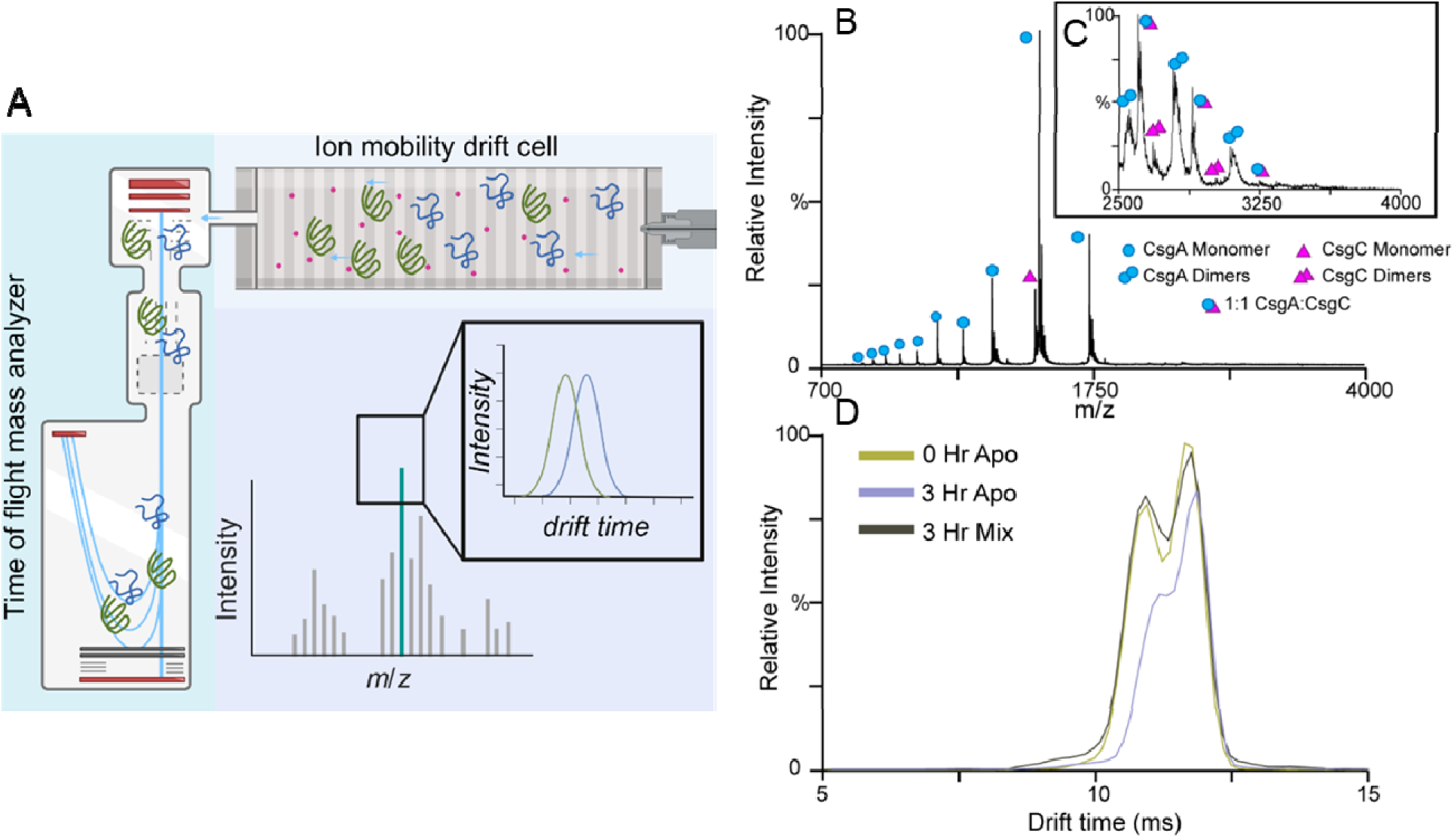
IM-MS reveals a 1:1 CsgA and CsgC heterodimer complex in solution and the stabilization of intrinsically disordered CsgA monomers. A) Proteins were passed through an ion mobility drift cell that separates based on size. Next, proteins pass through a time-of-flight mass analyzer. In combination, IM-MS can provide three modes of information for a given ion: mass to charge ratio, drift time, and the intensity of species with those properties. B) Mass spectra for CsgA incubated with CsgC in a 1:1 molar ratio at 37 °C. Monomeric CsgA (blue circles), dimeric CsgA (blue double circles), monomeric CsgC (purple triangles), dimeric CsgC (double purple triangles) and 1:1 CsgA:CsgC complexes (blue circle and purple triangle) are annotated. A magnified MS spectrum C) showed that the complex is seen flanked on either side by dimeric CsgA and dimeric CsgC. D) Arrival time distribution of the 11+ monomer of CsgA in three experimental conditions: Apo CsgA in solution at 0 hr (olive trace), apo CsgA in solution after 3 hr (light purple trace), and CsgA in solution with CsgC after 3 hr (dark grey trace). Traces are overlapped to show the changes in ATD after 3 hr of incubation.

We then analyzed the IM arrival time distributions (ATDs) to assess conformational changes within CsgA and CsgC over time. The ATD data contains information regarding both ion shape and the number of conformer families present for a single protein ion population (36). Data shown in **Fig. 5D** displays three overlaid ATD profiles for CsgA 11+ monomer ions recorded after 0 and 3 hour incubations. In addition, similar IM ATD data is shown for CsgA/CsgC mixtures following a 3 hour incubation. All three datasets yield a bimodal ATD profile, consisting of both a more compacted (shorter IM drift times) and a more extended conformational family (longer IM drift times). A comparison of CsgA IM data collected at both 0 and 3 hour time points revealed the peak corresponding to the more compact conformation decreased significantly in intensity, resulting in overwhelmingly a more extended protein ion population. Overall, our IM ATD data suggested that the conformational ensemble of CsgA becomes more extended as protein aggregation progresses, an observation common to other amyloid proteins (37, 38). Importantly, when we compare our IM ATD data recorded for CsgA at 0 hour with ATDs recorded for CsgA/CsgC mixtures following 3 hours of incubation, we detected few significant differences, indicating that the initial, intrinsically disordered state of CsgA is better preserved in the presence of CsgC. The results discussed above were consistent across multiple CsgA charge states (**Fig. S9C and D**).

## Discussion

Amyloid formation can lead to a wide range of pathologies. Therefore, it is of great interest to discover and characterize molecules that can prevent pathogenic amyloid formation. Synthetic amyloid inhibitor peptides occupy a large biochemical space, from β-blocking D-amino acid peptides (39) to camelid nanobodies (40, 41). There is a short list of naturally occurring proteins which putatively inhibit functional amyloid formation. This list includes molecular chaperones and a diverse set of human and bacterial proteins which have similar 3D structures (18).

However, CsgC differs from the rest of the known amyloid inhibitor proteins in one significant way: CsgC remains the only example of an amyloid inhibitor protein that exists exclusively to moderate the amyloid formation of specific proteins. CsgC is part of the CsgBAC operon, therefore, when amyloid proteins CsgA and CsgB are expressed, the CsgC protein will also be expressed. Studying CsgC provides a unique opportunity to answer questions about how cells prevent amyloid formation and utilize functional amyloids, as well as how other structurally similar amyloid inhibitors may function.

CsgC is a highly effective anti-amyloid chaperone protein (17), however, CsgC’s mechanism of action has remained unclear. Early work found that CsgC could inhibit CsgA without the need for a hydrolysable energy source and at a low sub-stoichiometric ratio (17). Key to understanding the mechanism of CsgA amyloid inhibition is the nature of the CsgC interacting partner. It has previously been suggested that CsgC interacts with elongating CsgA fibers (42), and there are other amyloid inhibitor proteins that act to inhibit monomer addition to growing fiber ends (43). However, molecular chaperones that interfere with amyloid formation have been observed to interact with a variety of binding partners and inhibit aggregation at different steps along the amyloid formation process (44). Given the multiplex role of monomers during amyloid formation, interactions between an inhibitor and an amyloid-competent monomer results in a decrease in the rate of several steps along amyloid formation (44). Our current data is consistent with an interaction between CsgC and a monomeric, prefibrillar species of CsgA as well as fibrillar CsgA at an extremely sub-stoichiometric ratio.

### CsgC extends CsgA Lag Phase

When proteins self-assemble into amyloid fibers through a nucleation-dependent process they typically display sigmoidal kinetics (24). There are three phases to polymerization beginning with the so called “lag phase”, a rate-limiting period of protein folding and initial oligomerization (24). After the lag phase, a critical mass of nuclei has formed, and aggregation increases exponentially as monomers quickly add onto fiber ends and new fibers continue to sprout from nuclei. CsgA is known to act mostly through a primary nucleation pathway and not through secondary nucleation where new fibers sprout from the lateral edge of an existing fiber (18, 45, 46).

CsgA requires a longer time to begin rapid amyloid formation when CsgC is present in solution. We saw lag phase extension in almost every scenario we tested, which included four CsgC homologs in a wide range of substrate:CsgC concentrations (**Fig. 1**). The CsgC homologs we tested share between 68% and 33% sequence identity with *E. coli* CsgC and yet, they all affect CsgA in a similar manner. Moreover, there is a distinct correlation between the concentration of the CsgC proteins and how much the lag phase was extended (**Fig. 1**). This would indicate a true dose-response relationship between CsgC proteins and the extension of CsgA lag phase and thereby the extent of primary nucleation. If CsgC is affecting nuclei formation, CsgC could be interacting with a prenuclear species of CsgA.

### CsgC inhibits new nuclei formation and fibril elongation during ThT elongation phase

CsgC remains an active amyloid inhibitor during the rapid fiber formation phase of aggregation. We show two similar experiments in **Fig. 2** where CsgC was presented to CsgA during the elongation phase. In the first experiment, CsgC was added to CsgA at different times throughout aggregation and we monitored the effect of CsgC addition on the increase in ThT fluorescence (**Fig. 2A**). CsgA alone takes approximately 1.3 hours to transition from the lag to the elongation phase (**Fig. 2A**). When CsgC is added at time 0, we see little to no increase in ThT fluorescence throughout the time course of the experiment, consistent with total inhibition of nuclei formation (**Fig. 2A**). For all conditions tested where CsgC was added after time 0 there was a linear increase in ThT signal, until the signal eventually reached a plateau and flattened out (**Fig. 2A**). This indicates that the addition of CsgC after initial formation of CsgA nuclei yields partial inhibition of CsgA amyloid formation. Linear or isodesmic polymerization is consistent with a model of aggregation that is independent of nucleation (47). Nucleation-dependent polymerization requires the formation of high energy nuclei, a bottleneck for polymerization, followed by the very favorable addition of monomers to a growing fiber (48). CsgC-inhibited CsgA could present an isodesmic polymerization profile because the formation of new nuclei is being hindered while CsgA unfolded monomers can still add to fiber ends. CsgC could be fighting the early process of CsgA transitioning between the disordered-denatured state and disordered-collapsed state as described by Frieden (47). However, this result can also arise due to inhibition of new nuclei formation as well as inhibition of fibril elongation.

To determine whether CsgC inhibition is solely based on inhibition of nuclei formation, a similar experiment was performed in the presence of seeds to rescue CsgA amyloid formation. Adding preformed fiber seeds to monomeric CsgA provides a template for the monomers to quickly adopt an aggregation prone state and therefore add onto a growing fiber end. Seeds act as catalysts, effectively lowering the free energy requirements to fibril formation. A linear increase in ThT signal can be seen when CsgC is added to a seeded CsgA aggregation reaction (**Fig. 2B**) indicating that the addition of seeds does not recapitulate un-inhibited CsgA amyloid formation. Therefore, CsgC may be acting to inhibit nuclei formation as well as fibril elongation.

The surface plasmon resonance findings indicate robust binding between CsgC monomer and CsgA fibrils (**Fig. 3**). Additionally, Amylofit data fitted to CsgA ThT binding assays in the presence of varying concentrations of CsgC homologs indicate that addition of CsgC and CsgC homologs drastically effect the rate of fibril elongation (**Fig. S2-5**). Taken together, it is likely that CsgC inhibits at later stages in amyloid formation through a stable interaction with fibrils, sterically preventing addition of CsgA monomers for fibril elongation.

It is worth noting that a “secondary-nucleation dominant” model was chosen to fit the experimental data analyzed by Amylofit. Secondary nucleation is less prevalent in functional amyloids as compared to disease associated amyloids and has not been observed in wild-type and mutant CsgA fibril formation (46, 49). Despite this, CsgA aggregation best resembles an aggregation model that accounts for both fibril growth and a form of self-replication (50). Future directions include single-molecule imaging of CsgA amyloid fibril growth to quantify individual fibril growth rates in the presence or absence of CsgC (51).

### CsgC Transiently Interacts with the CsgA Monomer

While CsgC may inhibit CsgA fibril growth, its function *in vivo* is likely to prevent any initiation of amyloid assembly which would be cytotoxic to the cell. Therefore, we sought to determine the mechanism of CsgC inhibition of pre-nuclear CsgA. Surface plasmon resonance indicated no measurable binding between CsgC monomers and CsgA monomers (**Fig. 3B).**

However, after quickly passing through a CsgC-linked column, freshly purified CsgA monomers require approximately 50 additional minutes to begin forming ThT positive species (**Fig. 4B** and **D**). The CsgA monomers were quickly pushed through the column by hand, therefore, any interaction with immobilized CsgC was over a matter of seconds. And yet, there was lasting impression on the CsgA, since aggregation was delayed (**Fig. 4D**). This suggests that a fast CsgC-CsgA monomer interaction delays CsgA monomers from adopting an aggregation-prone state.

Using IM-MS, we detected 1:1 CsgC and CsgA heterodimers, which indicates that CsgA and CsgC monomers can interact with each other (**Fig. 5B**). The heterodimer peaks were in low abundance, consistent with the concept of a transient interaction between CsgA and CsgC monomers. Indeed, the heterodimer peaks are the first direct observation of an interaction between full length CsgC and CsgA monomers.

### CsgC Maintains Intrinsically Disordered Monomeric CsgA

Much can be gleamed from focusing on the IM-MS signals associated with CsgA monomers. Multiple charge states of CsgA monomers can be seen in the mass spectrum, consistent with an intrinsically disordered protein (**Fig. 5B**). We analyzed how the CsgA monomer’s shape changed by examining the ATD over time (**Fig. 5D, S9C-D**). The ATD profile suggests that the conformation of CsgA became more extended as aggregation progresses, a change commonly observed with other amyloid proteins as well (**Fig. 5D, S9C-D**) (37, 38). Remarkably, the presence of CsgC in solution causes the ATD of CsgA monomers to remain consistent throughout the time course of our experiment (**Fig. 5D, S9C-D**). Since CsgA monomers start as intrinsically disordered, CsgC must be pushing CsgA monomers away from the aggregation-prone state by promoting or stabilizing CsgA’s intrinsically disordered fold. This is consistent with the findings from **Fig. 4**, illustrating that transient interaction between CsgA and CsgC leads to a memorable but temporary inhibition of CsgA amyloid formation.

## Conclusions

We utilized an array of biophysical methods to show that CsgC interacts transiently with an intrinsically disordered, monomeric CsgA to disrupt new nuclei formation in an ATP-independent fashion. This model of inhibition fits within the greater context of curli biogenesis:

CsgC is a periplasmic protein tasked with inhibiting amyloid formation by CsgA within the intermembrane space to prevent the formation of cytotoxic intracellular aggregates while also allowing for efficient secretion of monomeric and intrinsically disordered CsgA for subsequent amyloid formation on the bacterial surface. We also show that CsgC is capable of inhibiting CsgA fibril formation even in the presence of CsgA nuclei, likely through stably interacting with the amyloid fibril preventing further growth. Additionally, we show that CsgC utilizes a non-canonical chaperone mechanism by interacting with its client protein transiently, remodeling protein structure independently of a hydrolysable energy source, and promoting the unstructured fold state. We propose that CsgC, and other similarly structured amyloid inhibitors, may represent a new class of chaperone proteins capable of potent protein remodeling activity.

## Data availability

All data generated or analyzed during this study are included in this article or are available from the corresponding author upon reasonable request.

## Supporting information

This article contains supporting information.

## Acknowledgments

We thank members of the Chapman, Olofsson, and Ruotolo lab for their helpful suggestions and discussions. We also thank Georg Meisl for advice on using the Amylofit program correctly.

## Author Contributions

AB, DK, MC contributed conception and design of the study. AB, DK, YH, SJ, DL, LK, EG, and RA performed the experiments. AO and BR provided materials and expertise. AB and DK wrote the manuscript; AB, DK, and MC contributed to editing and revising the manuscript.

## Funding and additional information

This work was supported by the National Institutes of Health grant no. R01 GM118651 and R21 AI137535 to M.R.C., R01 GM095832 to B.R., and NIH T32 AI007528 to DK.

## Conflict of Interest

The authors declare that they have no conflicts of interest with the contents of this article.

## Abbreviations

SPR: Surface Plasmon Resonance
SDS-PAGE: sodium dodecyl sulphate-polyacrylamide gel electrophoresis
IM-MS: ion mobility-mass spectrometry
TOF: time-of-flight
ThT: Thioflavin-T
EC: *E. coli*
CY: *Citrobacter youngae*
CD: *Citrobacter youngae*
HA: *Hafnia alvei*
MRE: mean residual error
HFIP: hexafluoroisopropanol
CCS: collisional cross section
ATD: arrival time distribution.

## Supporting Information

**Fig. S1.**
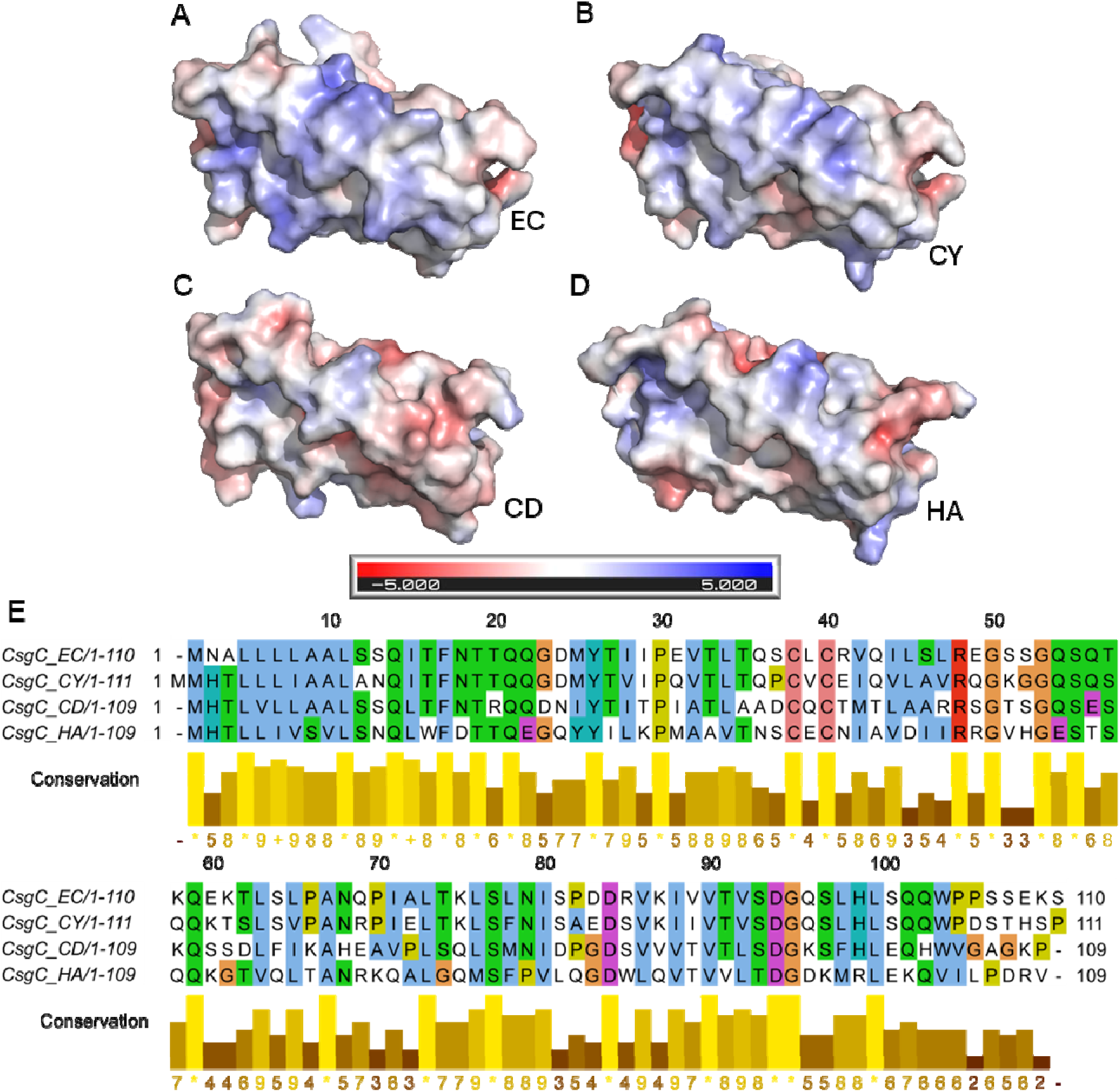
CsgC homolog electrostatic surface potential maps and sequence alignment. **A-D)** AlphaFold 2.0 predicted structures for CsgC EC, CY, CD, and HA were loaded into Pymol. The four proteins were aligned using the default align method. The APBS plugin (52) was used to display the electrostatic surface potential of the proteins and the automatically generated charge color guide is displayed. **E)** A multiple sequences alignment of the four CsgC homologs was performed using the MAFFT method (53). The amino acids are colored using the Clustal X color scheme. The conservation of physico-chemical properties of each position in the alignment was calculated and a numbered index was provided; a * denotes complete sequence identity (54).

**Fig. S2.**
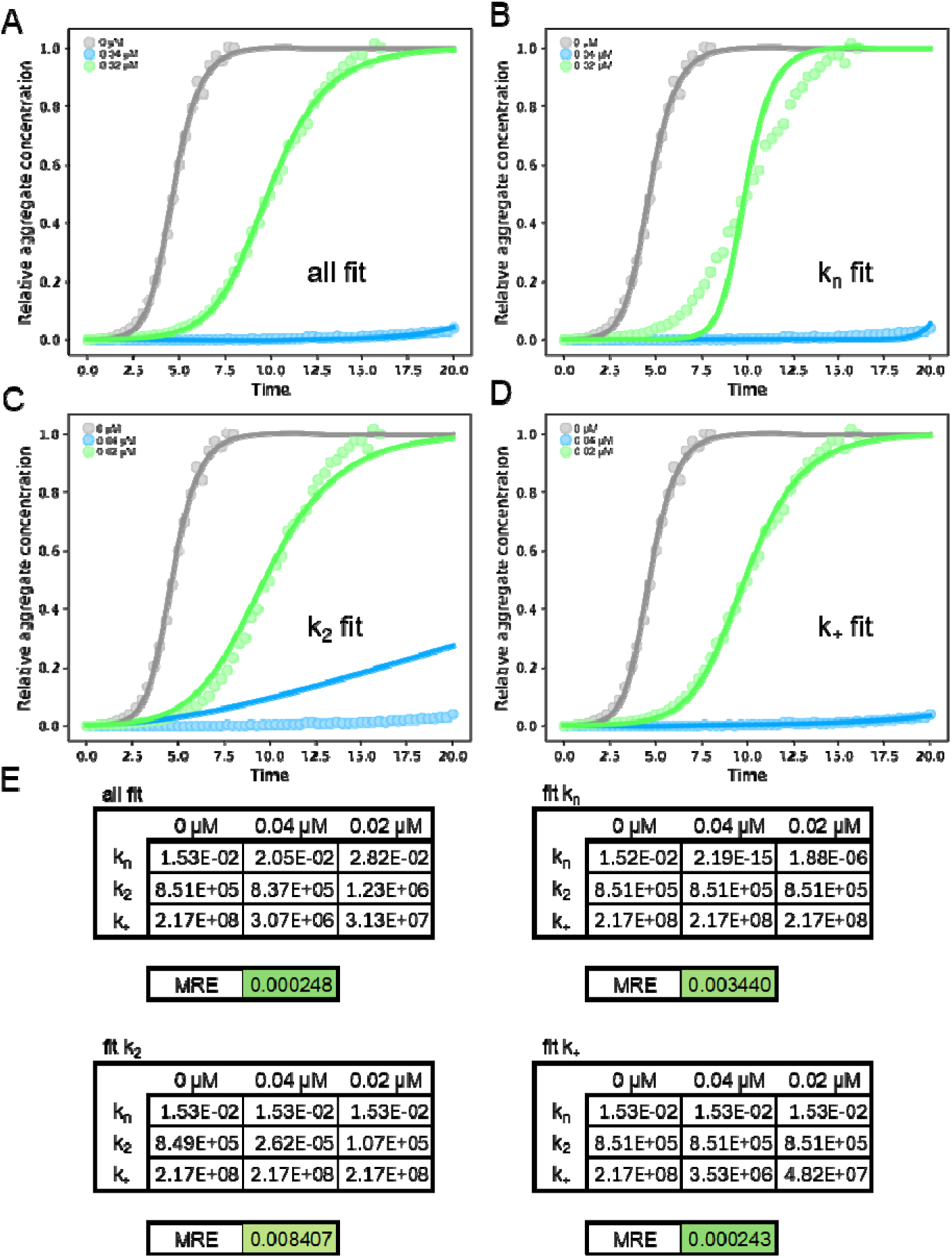
The effect of CsgC EC on CsgA amyloid formation kinetics. CsgA was freshly purified and a ThT binding assay was performed using 20 µM CsgA mixed with CsgC EC added to the labeled final concentration. The data was entered into Amylofit (27) and any curve which showed no increase in signal over the time course was removed. The secondary nucleation dominated model was used to determine the global or individual fit of each rate constant parameter. **A)** At first, all rate constant parameters were set to “fit” to individually determine their best fit values. **B-D)** Subsequently, all parameters but one were set to “global constant” and used the values determined in the individual fit analysis of uninhibited CsgA. The one remaining parameter was set to “fit” and the value determined in the individual fit analysis of uninhibited CsgA was used as the initial guess. **E)** By testing each rate constant in this manner, a single mean squared residual error (MRE) was produced when all inhibitor concentrations were fit globally.

**Fig. S3.**
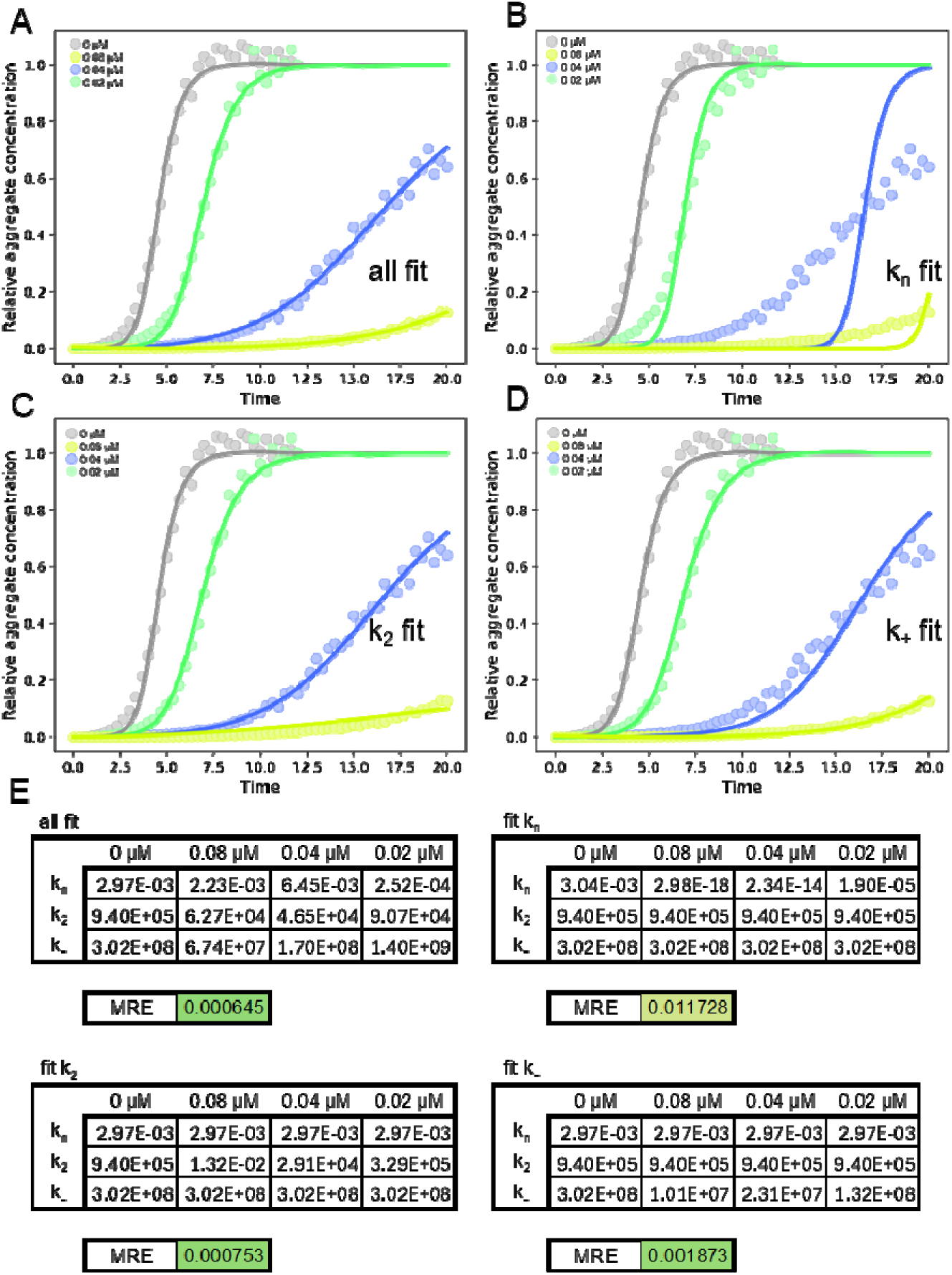
The effect of CsgC CY on CsgA amyloid formation kinetics. CsgA was freshly purified and a ThT binding assay was performed using 20 µM CsgA mixed with CsgC CY added to the labeled final concentration. The data was entered into Amylofit (27) and any curve which showed no increase in signal over the time course was removed. The secondary nucleation dominated model was used to determine the global or individual fit of each rate constant parameter. **A)** At first, all rate constant parameters were set to “fit” to individually determine their best fit values. **B-D)** Subsequently, all parameters but one were set to “global constant” and used the values determined in the individual fit analysis of uninhibited CsgA. The one remaining parameter was set to “fit” and the value determined in the individual fit analysis of uninhibited CsgA was used as the initial guess. **E)** By testing each rate constant in this manner, a single mean squared residual error (MRE) was produced when all inhibitor concentrations were fit globally.

**Fig. S4.**
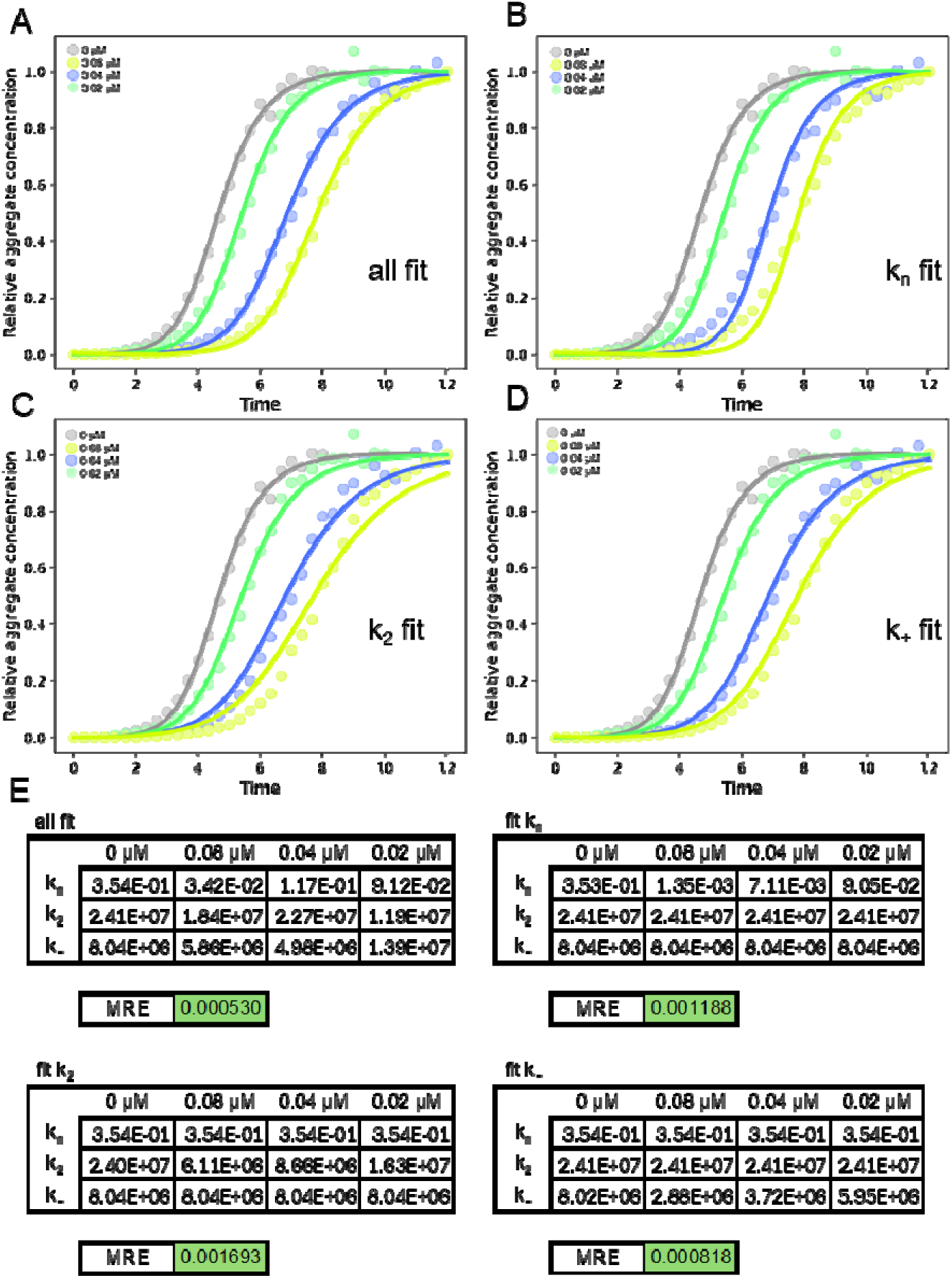
The effect of CsgC CD on CsgA amyloid formation kinetics. CsgA was freshly purified and a ThT binding assay was performed using 20 µM CsgA mixed with CsgC CD added to the labeled final concentration. The data was entered into Amylofit (27) and any curve which showed no increase in signal over the time course was removed. The secondary nucleation dominated model was used to determine the global or individual fit of each rate constant parameter. **A)** At first, all rate constant parameters were set to “fit” to individually determine their best fit values. **B-D)** Subsequently, all parameters but one were set to “global constant” and used the values determined in the individual fit analysis of uninhibited CsgA. The one remaining parameter was set to “fit” and the value determined in the individual fit analysis of uninhibited CsgA was used as the initial guess. **E)** By testing each rate constant in this manner, a single mean squared residual error (MRE) was produced when all inhibitor concentrations were fit globally.

**Fig. S5.**
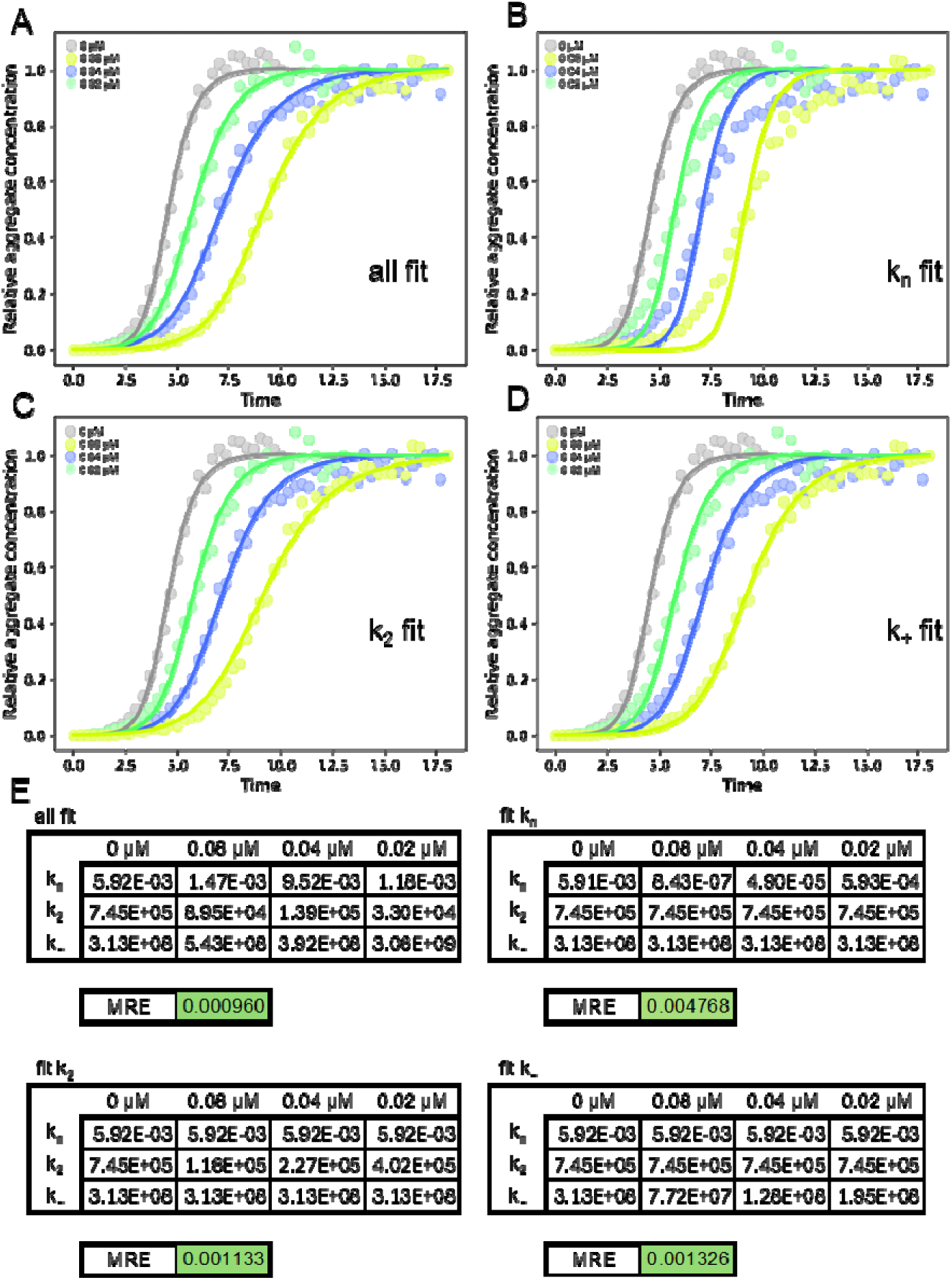
The effect of CsgC HA on CsgA amyloid formation kinetics. CsgA was freshly purified and a ThT binding assay was performed using 20 µM CsgA mixed with CsgC HA added to the labeled final concentration. The data was entered into Amylofit (27) and any curve which showed no increase in signal over the time course was removed. The secondary nucleation dominated model was used to determine the global or individual fit of each rate constant parameter. **A)** At first, all rate constant parameters were set to “fit” to individually determine their best fit values. **B-D)** Subsequently, all parameters but one were set to “global constant” and used the values determined in the individual fit analysis of uninhibited CsgA. The one remaining parameter was set to “fit” and the value determined in the individual fit analysis of uninhibited CsgA was used as the initial guess. **E)** By testing each rate constant in this manner, a single mean squared residual error (MRE) was produced when all inhibitor concentrations were fit globally.

**Fig. S6.**
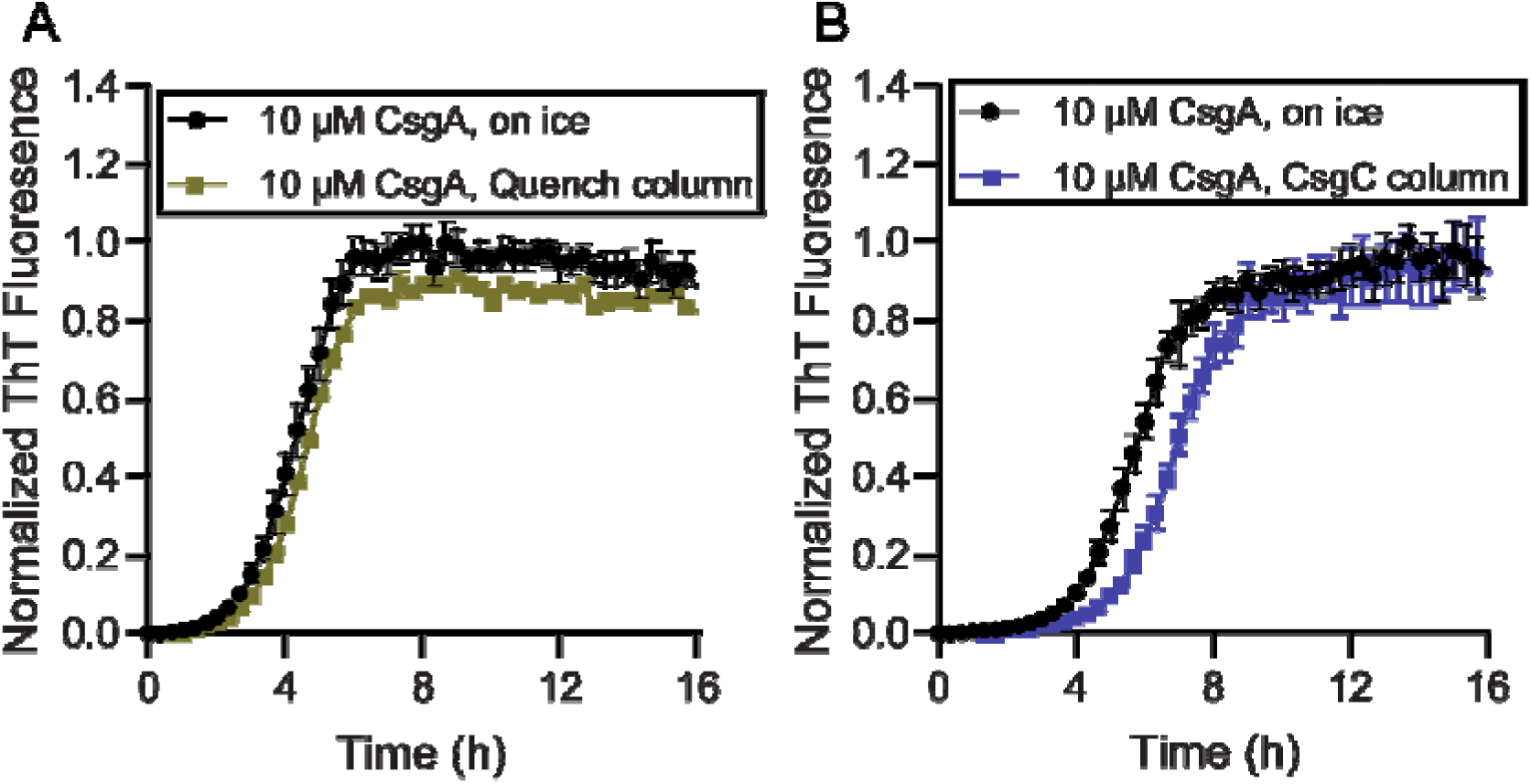
Additional Pull Down Assay Data. ThT binding assays of several controls for the modified NHS-resin pull down assays. CsgA was reserved on ice or passed through a column containing **A)** Tris-linked and quenched resin or **B)** CsgC-linked resin. Eluent CsgA was then diluted to 10 µM in phosphate buffer for a ThT binding assay to observe amyloid aggregation kinetics. Tris-linked resin has little to no effect on CsgA aggregation kinetics while CsgC-linked resin delayed CsgA amyloid formation.

**Fig. S7.**
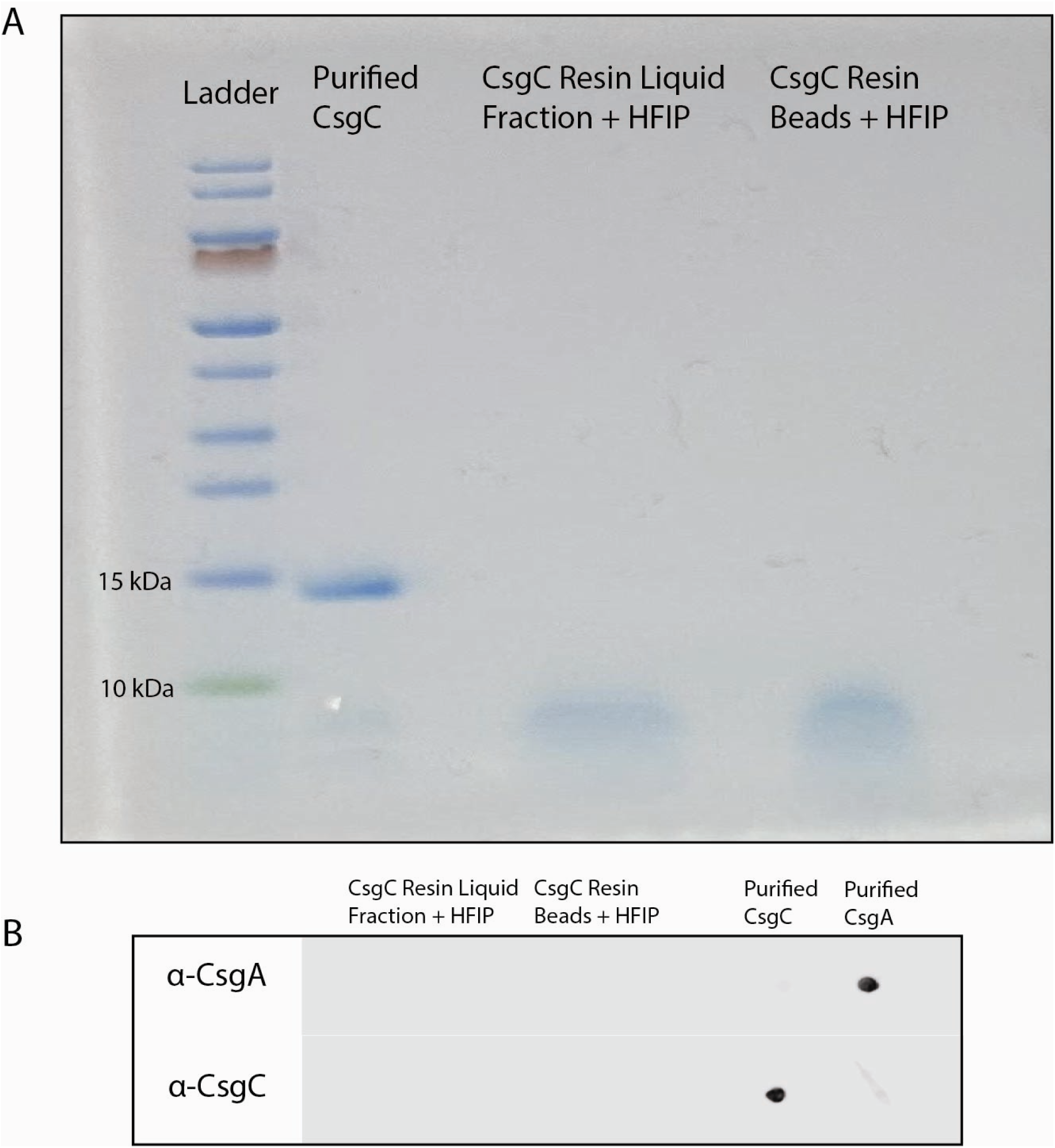
CsgA is not retained on the CsgC-linked resin. **A)** SDS-Page analysis of CsgC-linked resin that has interacted with CsgA. Resin was separated into a liquid fraction and a bead fraction and treated with HFIP. No bands were detected indicating that there is no CsgC or CsgA removed from the CsgC-linked beads. **B)** Dot blot of samples from (A) illustrates that there is no detectable CsgC or CsgA removed from the CsgC-linked beads following HFIP treatment.

**Fig. S8.**
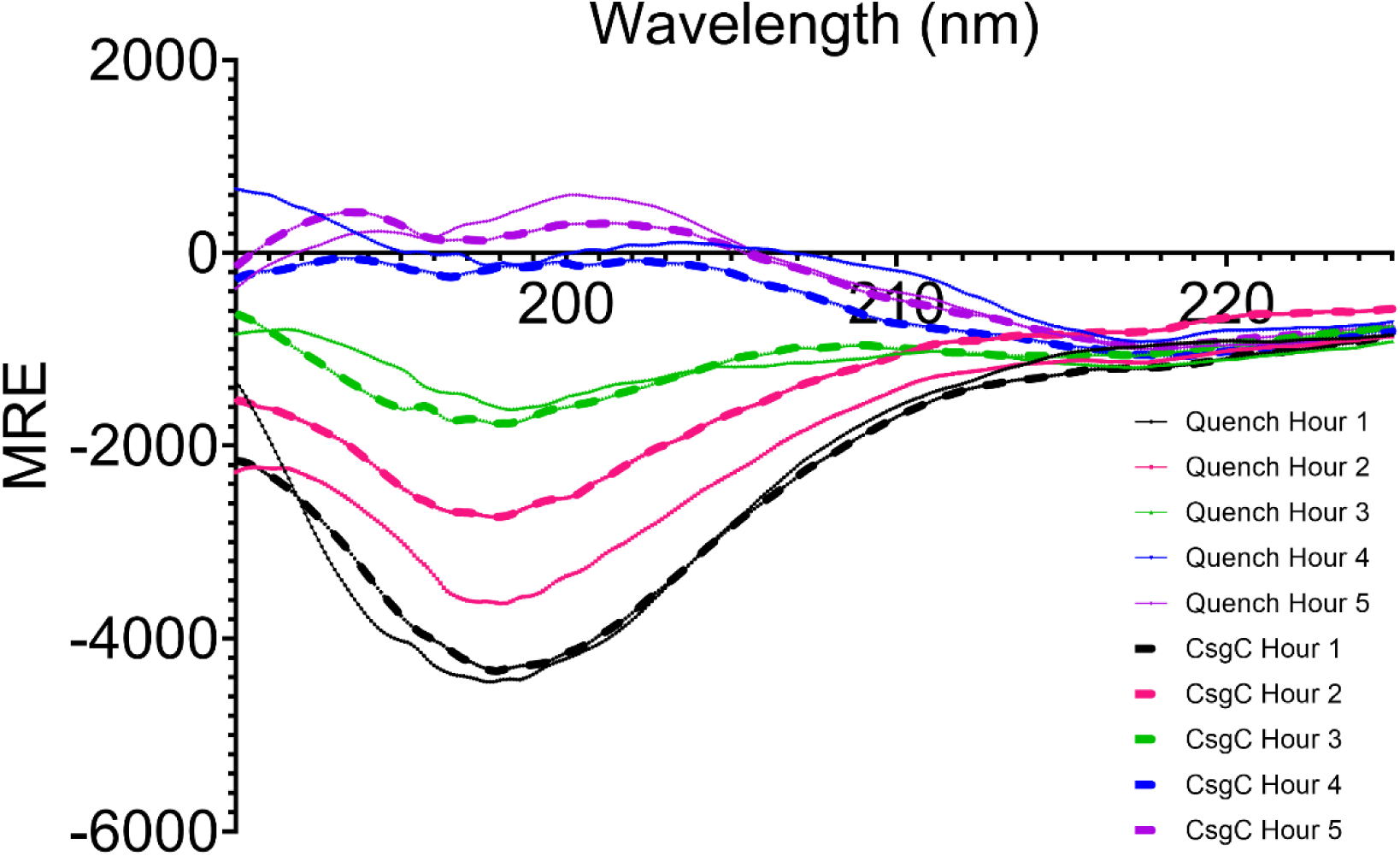
Circular Dichroism of CsgA eluted from quenched or CsgC-linked resin. Circular dichroism of 15 µM CsgA eluted from quenched NHS-Resin (called Quench Hour X, solid lines) or CsgC-linked resin (called CsgC Hour X, dashed lines) X hours following purification. Measurements were taken in triplicate and averaged. Measurements were also taken over two independent assays to account for sampling order.

**Fig. S9.**
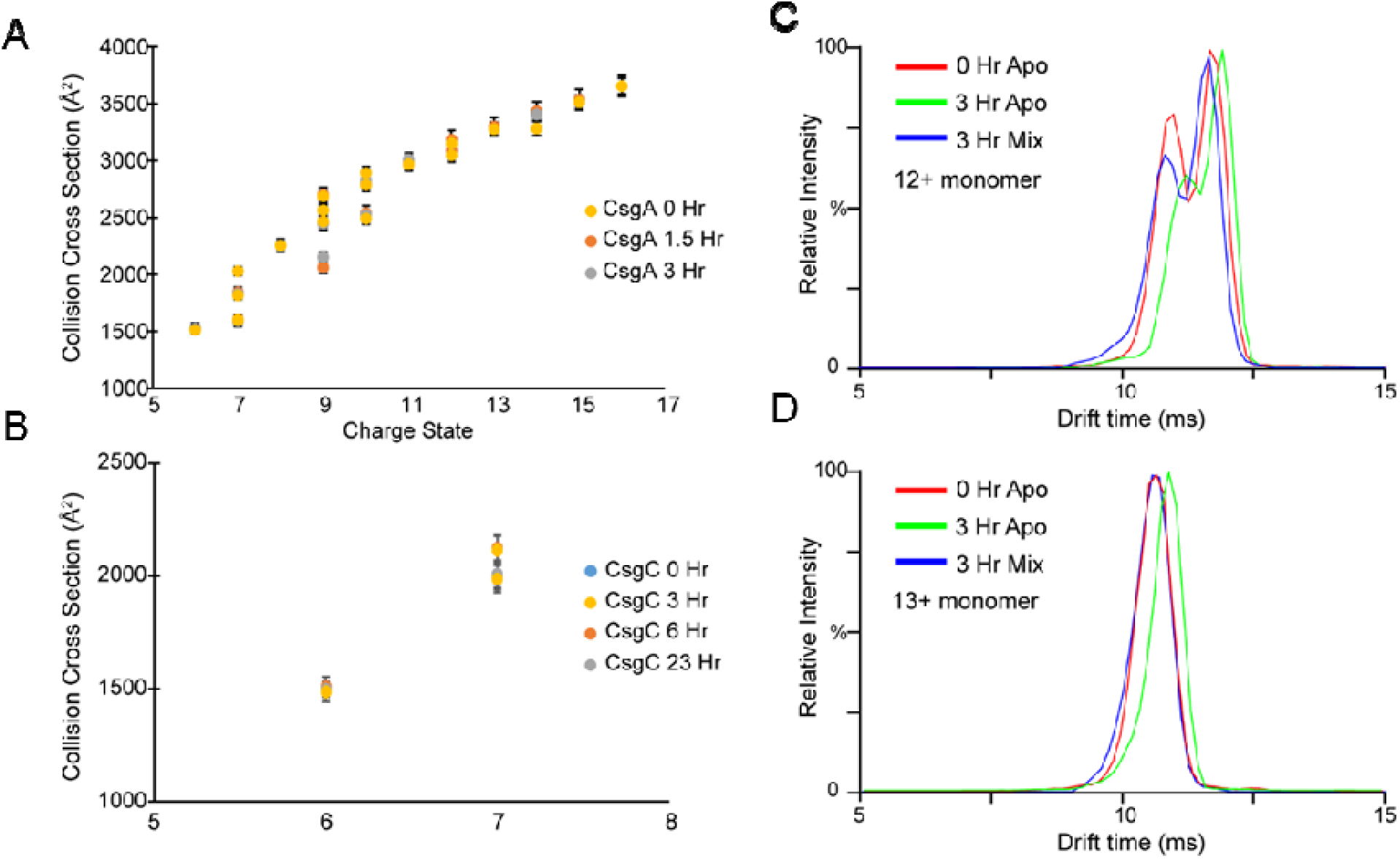
Additional IM-MS Data. CCS values derived from arrival time distribution of **A)** CsgA monomers and **B)** CsgC monomers. **C and D)** Arrival time distribution of the 12+ and 13+ monomer of CsgA in three experimental conditions: Apo CsgA in solution at 0 hr (red trace), apo CsgA in solution after 3 hr (green trace), and CsgA in solution with CsgC after 3 hr (blue trace). Traces are overlapped to show the changes in ATD after 3 hr of incubation.

## Supporting Tables

**Supporting Table 1.**
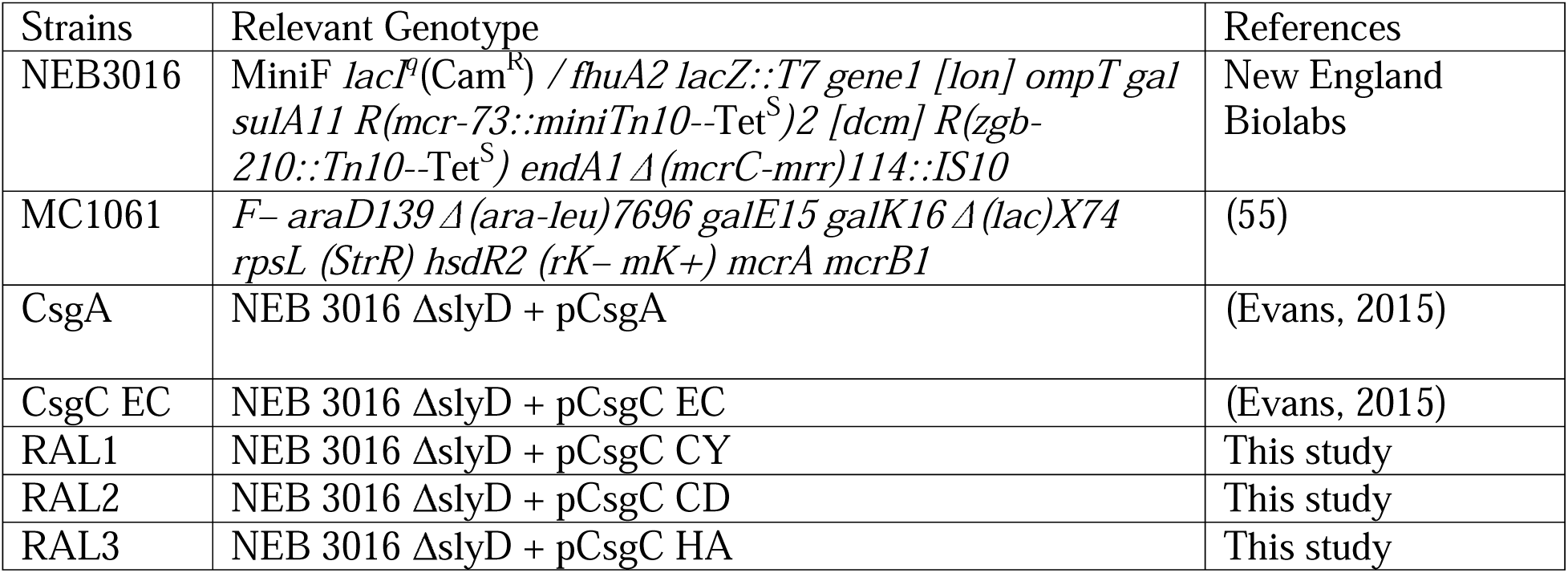
Strains used in this study.

**Supporting Table 2.**
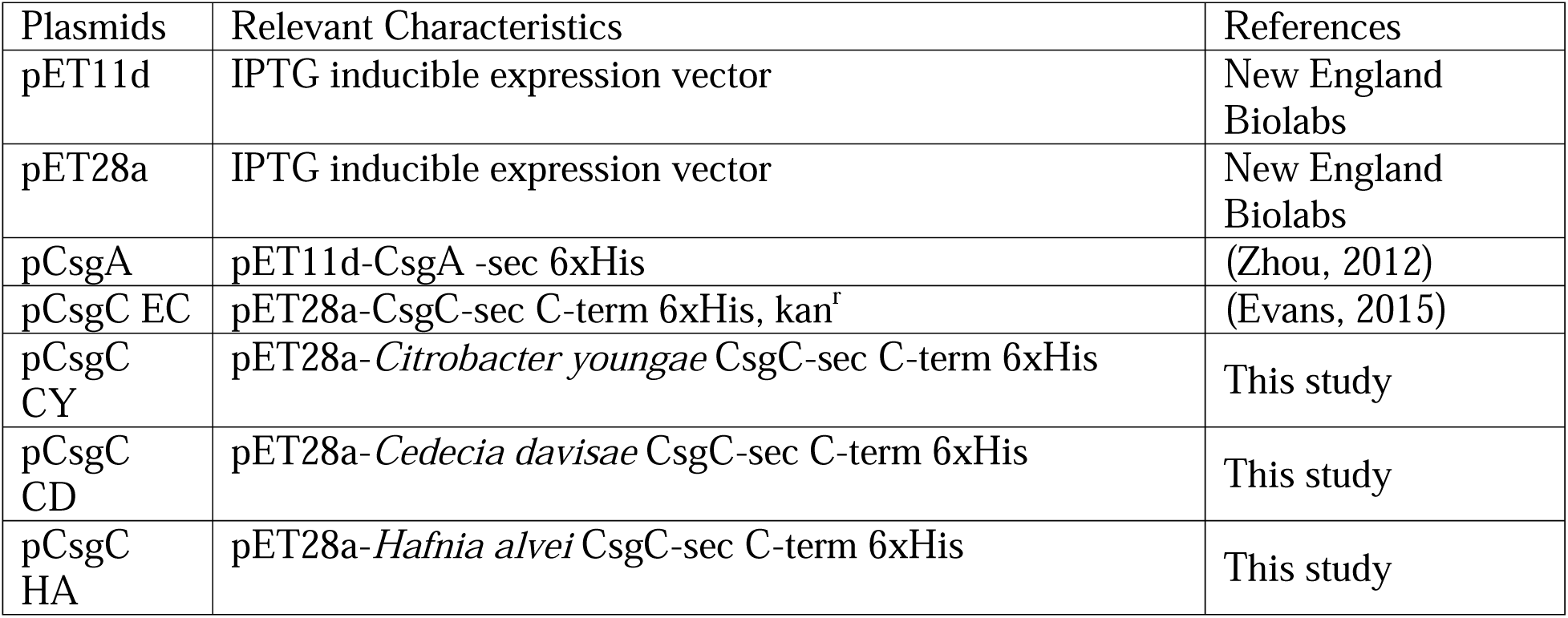
Plasmids used in this study.

**Supporting Table 3.**
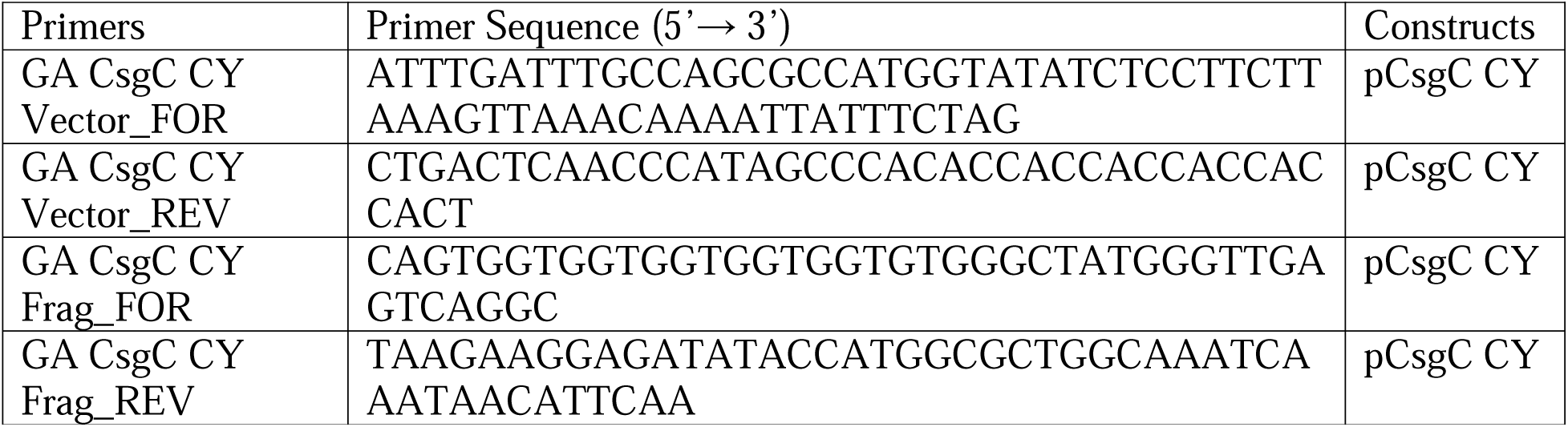
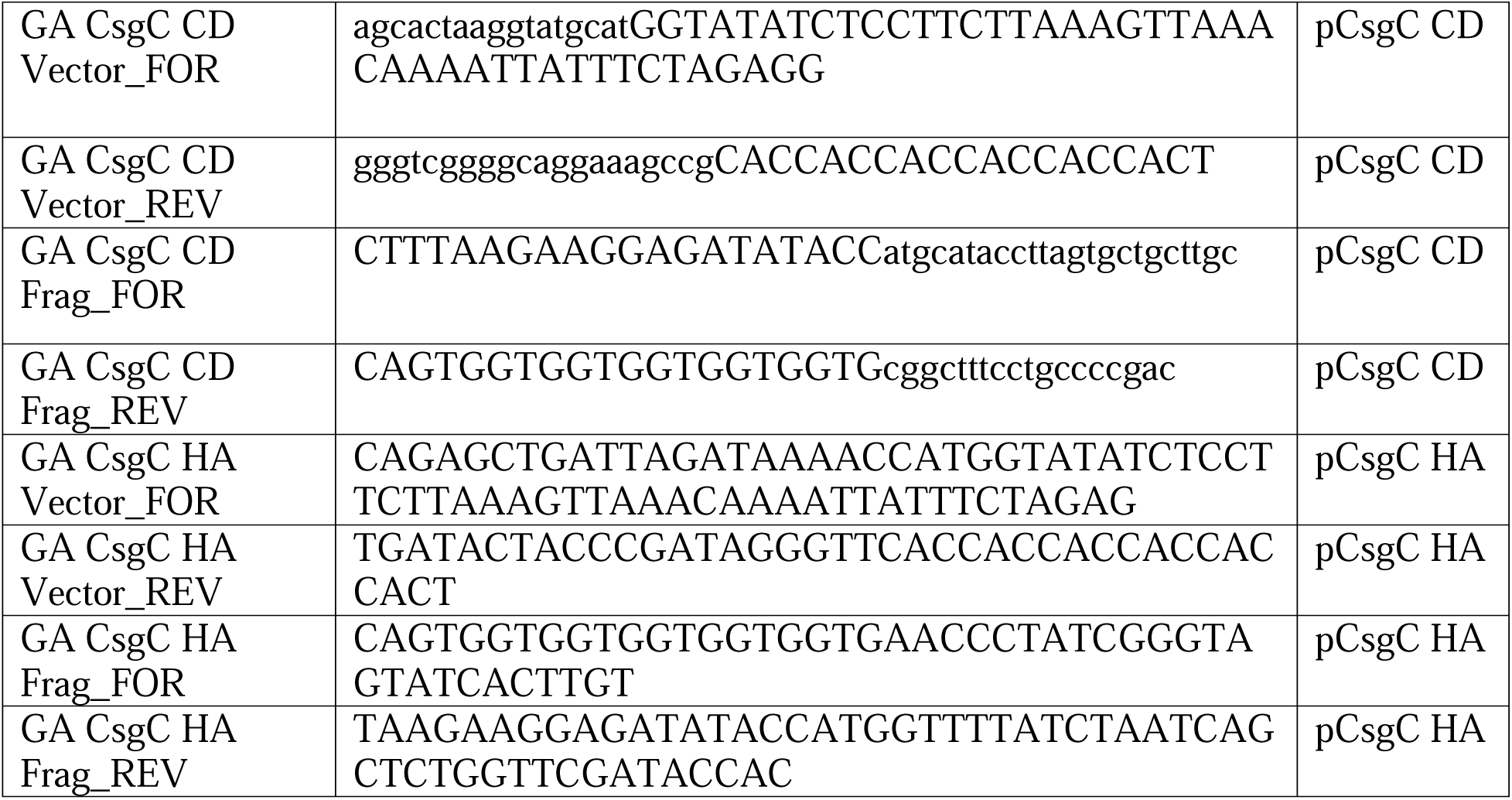
Primers used in this study.

## Notes

### Competing Interest Statement

The authors have declared no competing interest.

### Summary of Updates

Added recognition for co-first authorship. No other changes were made.

