## Supporting Information for "The bacterial chaperone CsgC inhibits functional amyloid CsgA formation by promoting the intrinsically disordered pre-nuclear state"

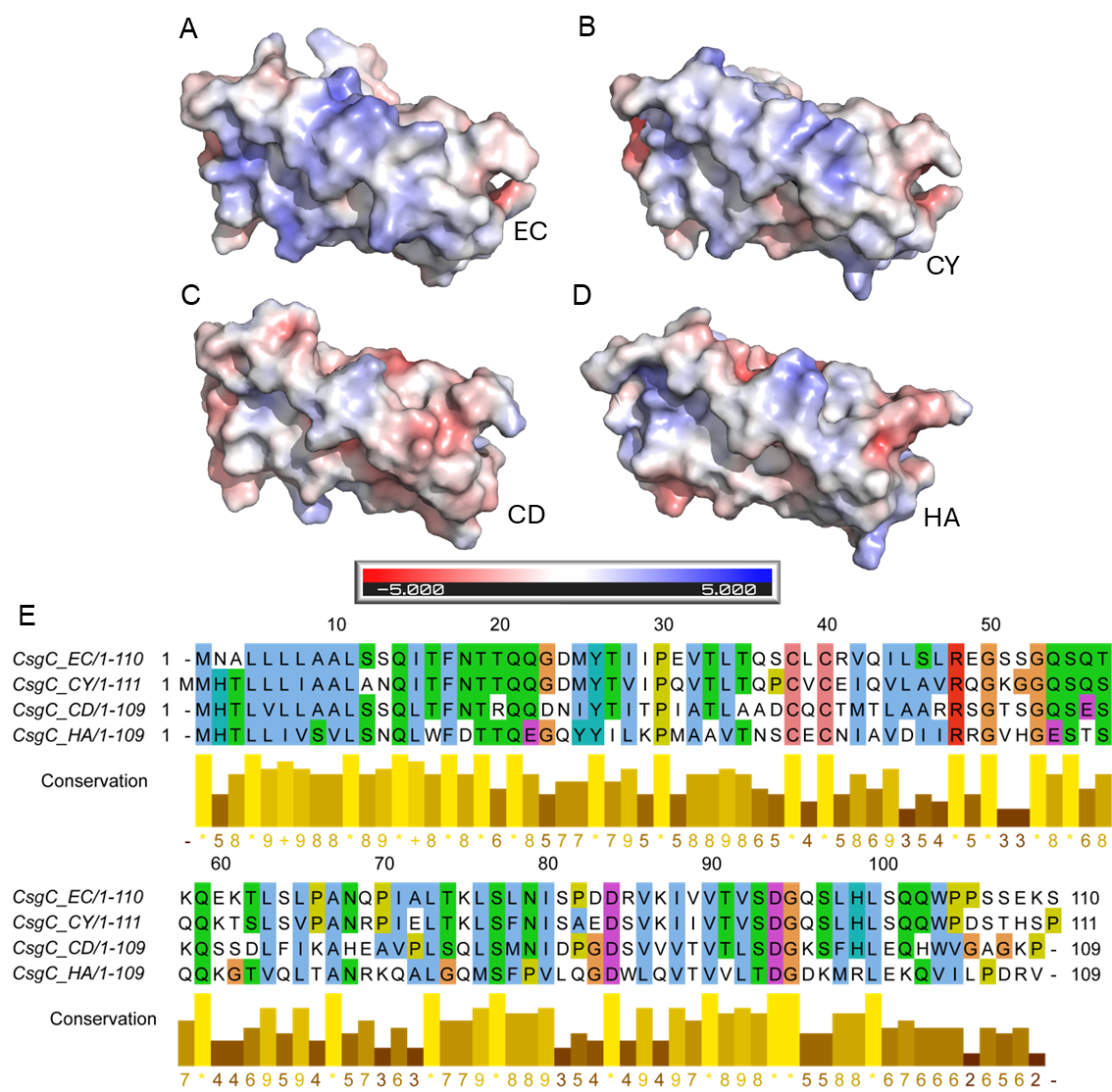

**Fig. S1. CsgC homolog electrostatic surface potential maps and sequence alignment.** **A-D)** AlphaFold 2.0 predicted structures for CsgC EC, CY, CD, and HA were loaded into Pymol. The four proteins were aligned using the default align method. The APBS plugin (52) was used to display the electrostatic surface potential of the proteins and the automatically generated charge color guide is displayed. **E)** A multiple sequences alignment of the four CsgC homologs was performed using the MAFFT method (53). The amino acids are colored using the Clustal X color scheme. The conservation of physico-chemical properties of each position in the alignment was calculated and a numbered index was provided; a * denotes complete sequence identity (54).

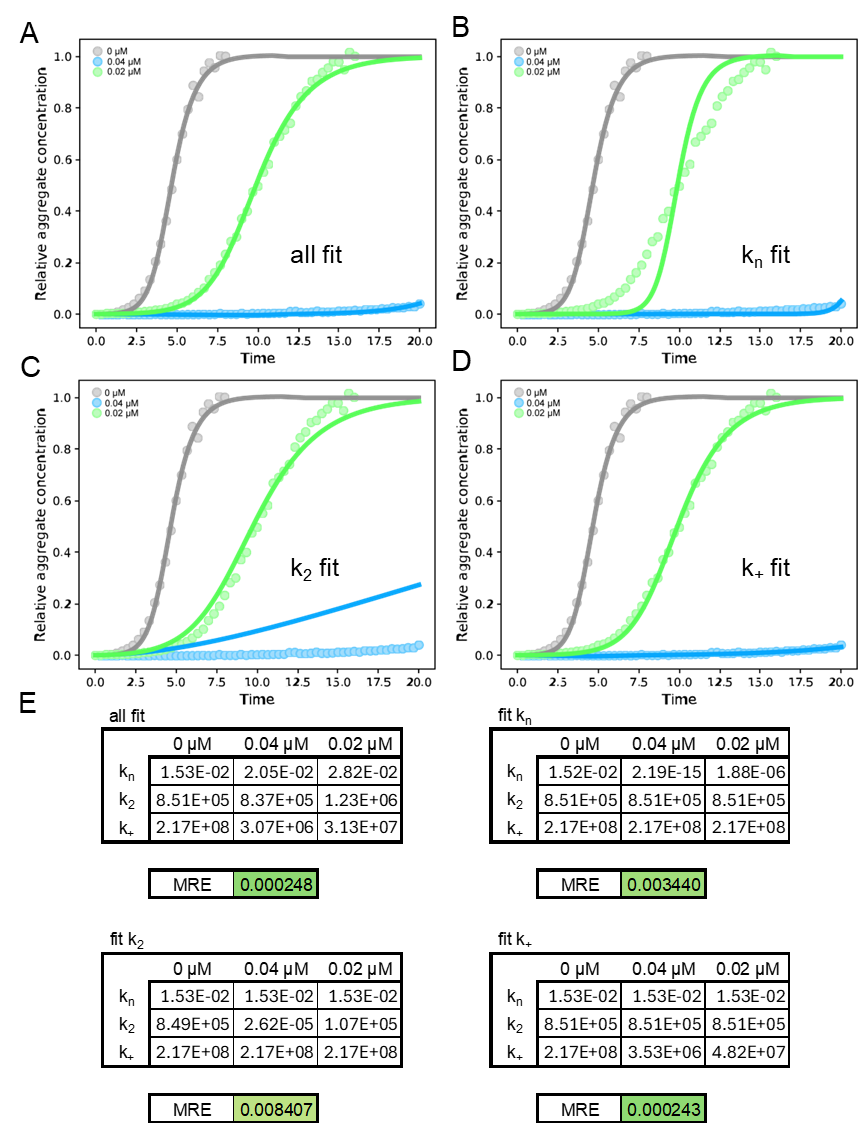

**Fig. S2. The effect of CsgC EC on CsgA amyloid formation kinetics.** CsgA was freshly purified and a ThT binding assay was performed using 20 µM CsgA mixed with CsgC EC added to the labeled final concentration. The data was entered into Amylofit (27) and any curve which showed no increase in signal over the time course was removed. The secondary nucleation dominated model was used to determine the global or individual fit of each rate constant parameter. **A)** At first, all rate constant parameters were set to “fit” to individually determine their best fit values. **B-D)** Subsequently, all parameters but one were set to “global constant” and used the values determined in the individual fit analysis of uninhibited CsgA. The one remaining parameter was set to “fit” and the value determined in the individual fit analysis of uninhibited CsgA was used as the initial guess. **E)** By testing each rate constant in this manner, a single mean squared residual error (MRE) was produced when all inhibitor concentrations were fit globally.

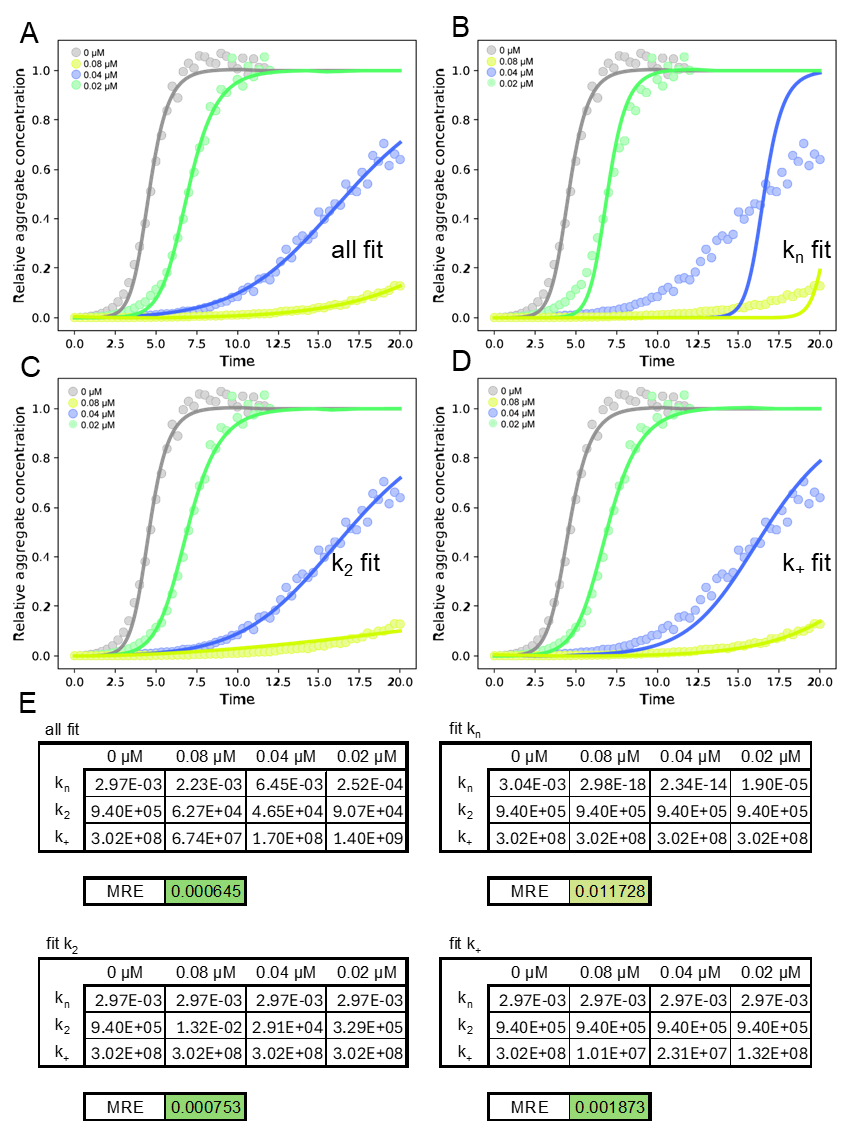

**Fig. S3. The effect of CsgC CY on CsgA amyloid formation kinetics.** CsgA was freshly purified and a ThT binding assay was performed using 20 µM CsgA mixed with CsgC CY added to the labeled final concentration. The data was entered into Amylofit (27) and any curve which showed no increase in signal over the time course was removed. The secondary nucleation dominated model was used to determine the global or individual fit of each rate constant parameter. **A)** At first, all rate constant parameters were set to “fit” to individually determine their best fit values. **B-D)** Subsequently, all parameters but one were set to “global constant” and used the values determined in the individual fit analysis of uninhibited CsgA. The one remaining parameter was set to “fit” and the value determined in the individual fit analysis of uninhibited CsgA was used as the initial guess. **E)** By testing each rate constant in this manner, a single mean squared residual error (MRE) was produced when all inhibitor concentrations were fit globally.

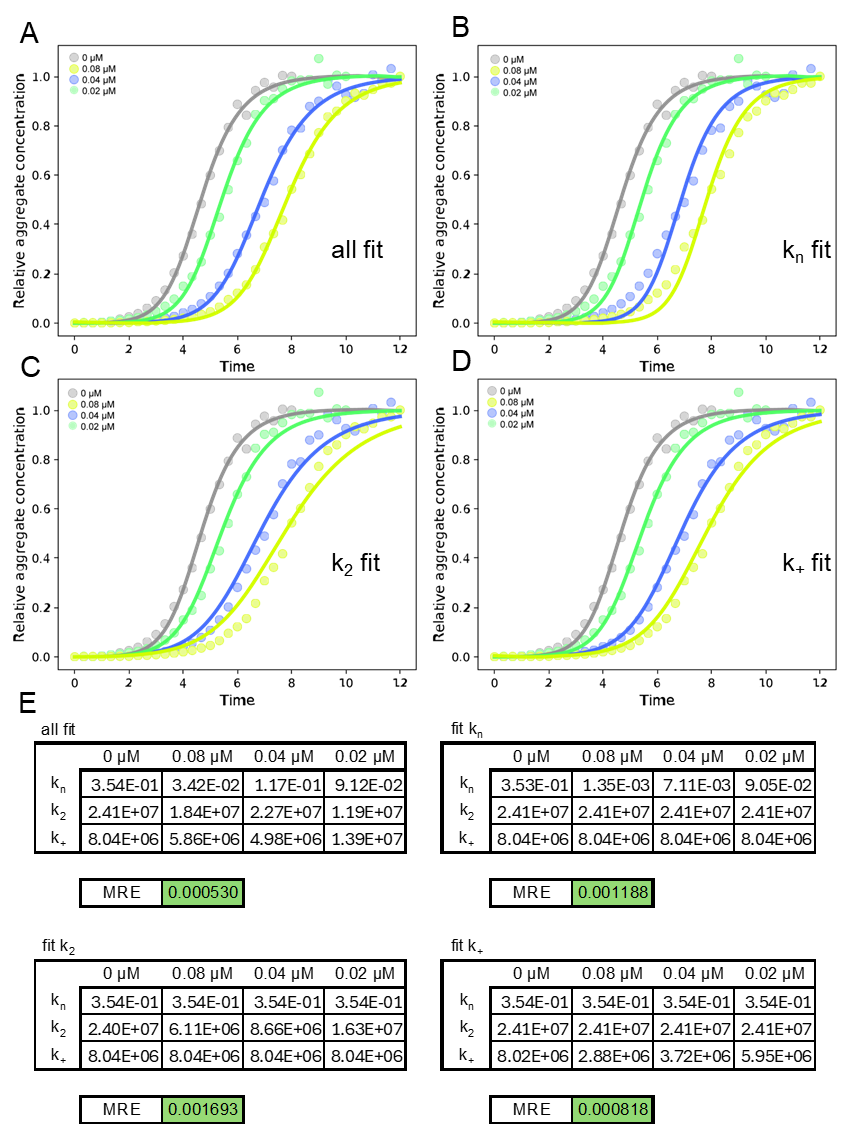

**Fig. S4. The effect of CsgC CD on CsgA amyloid formation kinetics.** CsgA was freshly purified and a ThT binding assay was performed using 20 µM CsgA mixed with CsgC CD added to the labeled final concentration. The data was entered into Amylofit (27) and any curve which showed no increase in signal over the time course was removed. The secondary nucleation dominated model was used to determine the global or individual fit of each rate constant parameter. **A)** At first, all rate constant parameters were set to “fit” to individually determine their best fit values. **B-D)** Subsequently, all parameters but one were set to “global constant” and used the values determined in the individual fit analysis of uninhibited CsgA. The one remaining parameter was set to “fit” and the value determined in the individual fit analysis of uninhibited CsgA was used as the initial guess. **E)** By testing each rate constant in this manner, a single mean squared residual error (MRE) was produced when all inhibitor concentrations were fit globally.

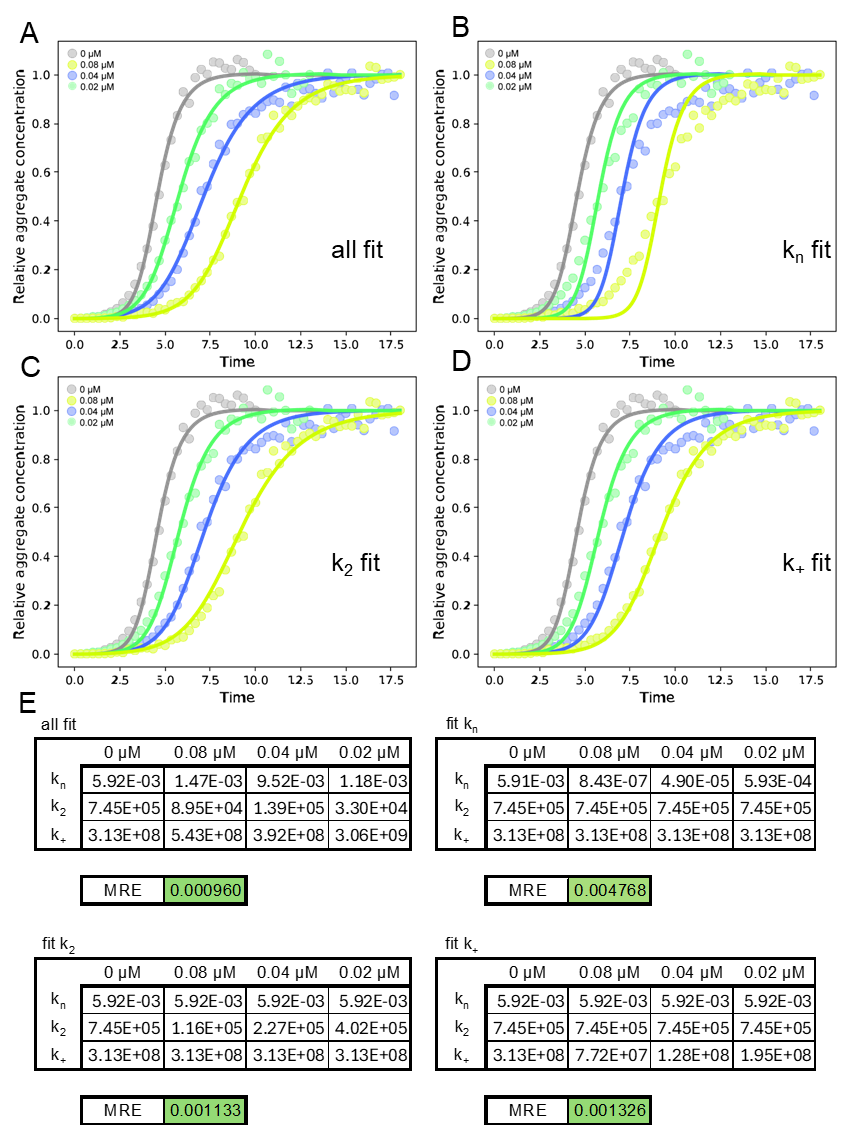

**Fig. S5. The effect of CsgC HA on CsgA amyloid formation kinetics.** CsgA was freshly purified and a ThT binding assay was performed using 20 µM CsgA mixed with CsgC HA added to the labeled final concentration. The data was entered into Amylofit (27) and any curve which showed no increase in signal over the time course was removed. The secondary nucleation dominated model was used to determine the global or individual fit of each rate constant parameter. **A)** At first, all rate constant parameters were set to “fit” to individually determine their best fit values. **B-D)** Subsequently, all parameters but one were set to “global constant” and used the values determined in the individual fit analysis of uninhibited CsgA. The one remaining parameter was set to “fit” and the value determined in the individual fit analysis of uninhibited CsgA was used as the initial guess. **E)** By testing each rate constant in this manner, a single mean squared residual error (MRE) was produced when all inhibitor concentrations were fit globally.

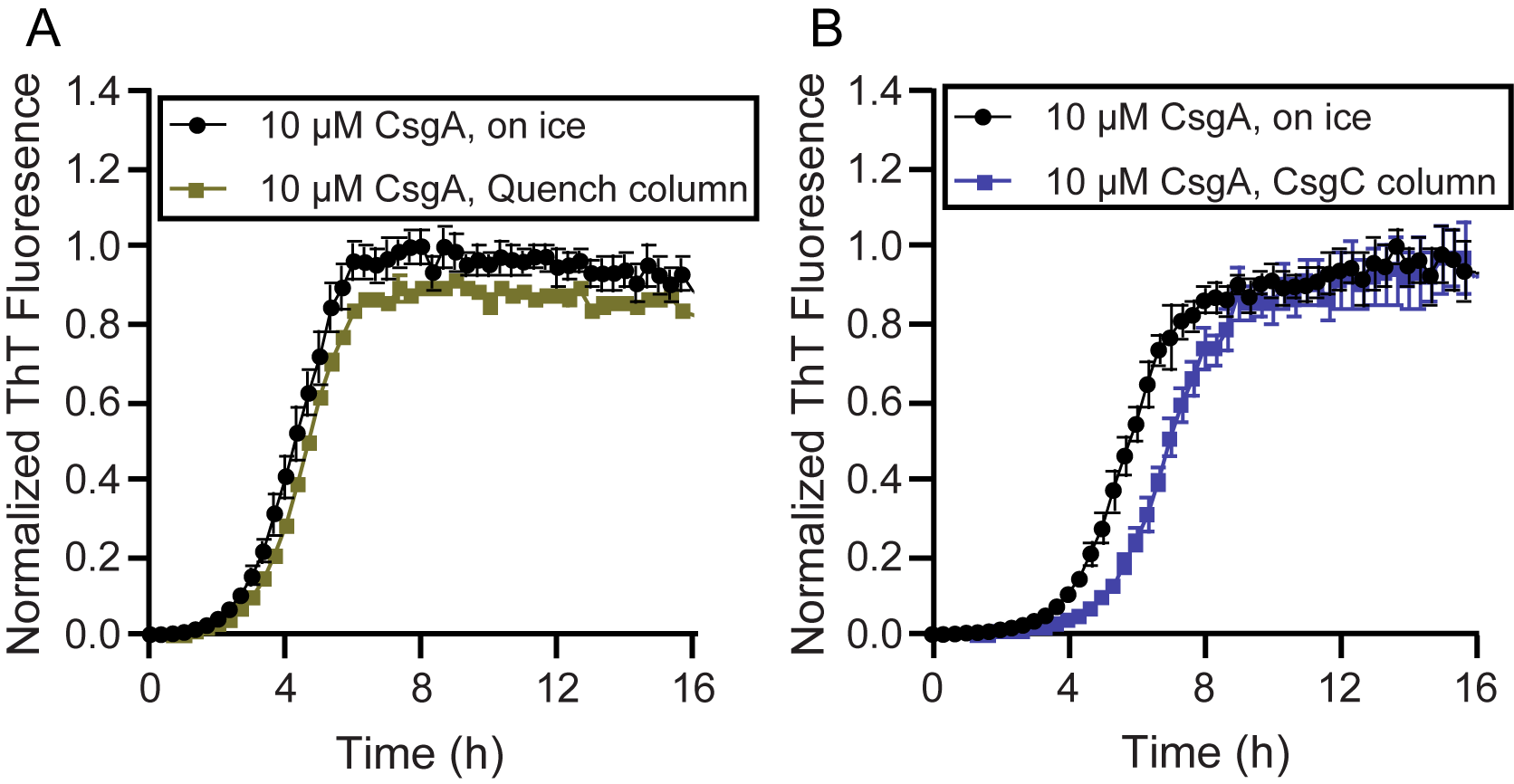

Fig. S6. Additional Pull Down Assay Data. ThT binding assays of several controls for the modified NHS-resin pull down assays. CsgA was reserved on ice or passed through a column containing A) Tris-linked and quenched resin or B) CsgC-linked resin. Eluent CsgA was then diluted to 10 µM in phosphate buffer for a ThT binding assay to observe amyloid aggregation kinetics. Tris-linked resin has little to no effect on CsgA aggregation kinetics while CsgC-linked resin delayed CsgA amyloid formation.

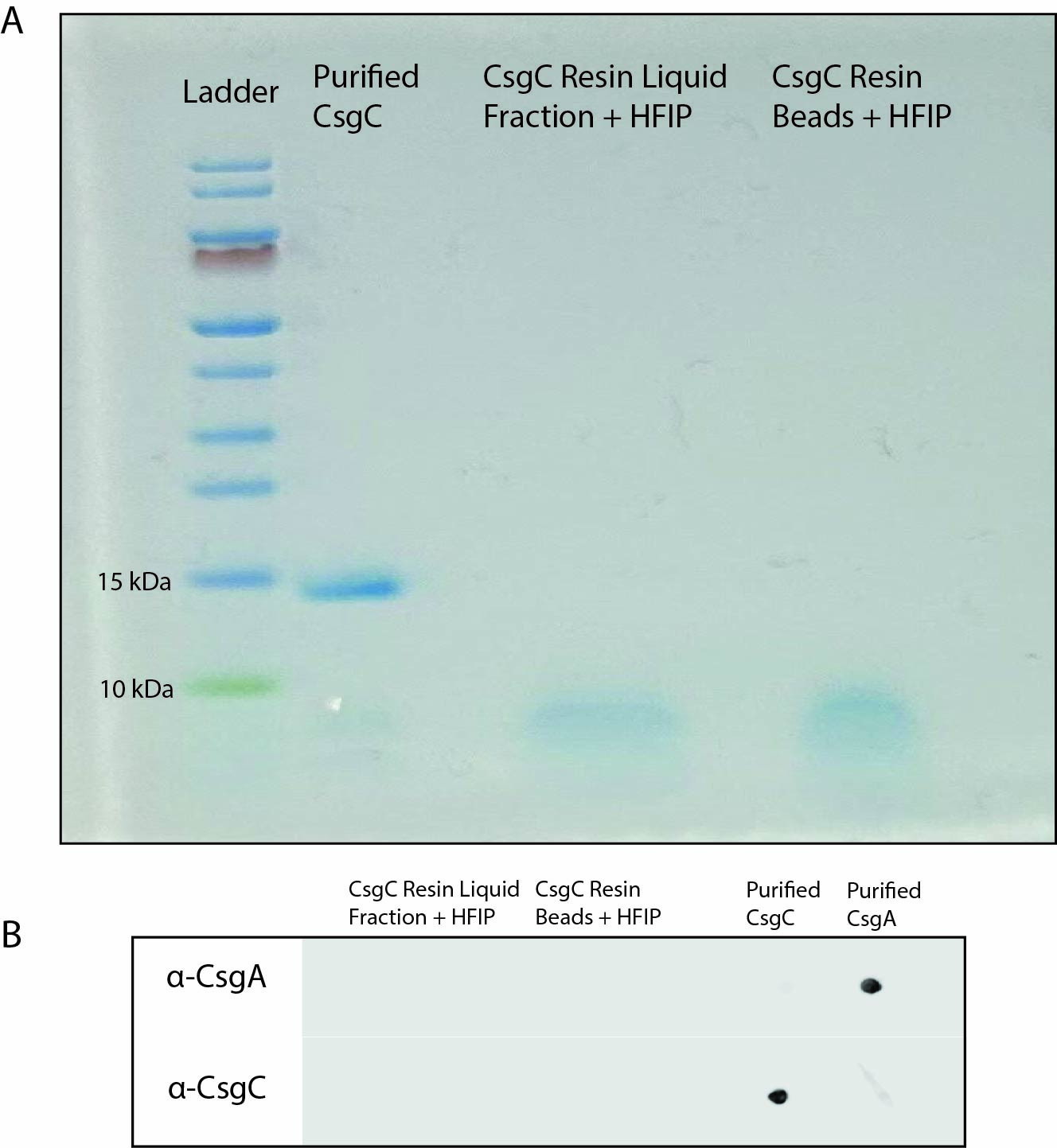

**Fig. S7. CsgA is not retained on the CsgC-linked resin. A)** SDS-Page analysis of CsgC-linked resin that has interacted with CsgA. Resin was separated into a liquid fraction and a bead fraction and treated with HFIP. No bands were detected indicating that there is no CsgC or CsgA removed from the CsgC-linked beads. **B)** Dot blot of samples from (A) illustrates that there is no detectable CsgC or CsgA removed from the CsgC-linked beads following HFIP treatment.

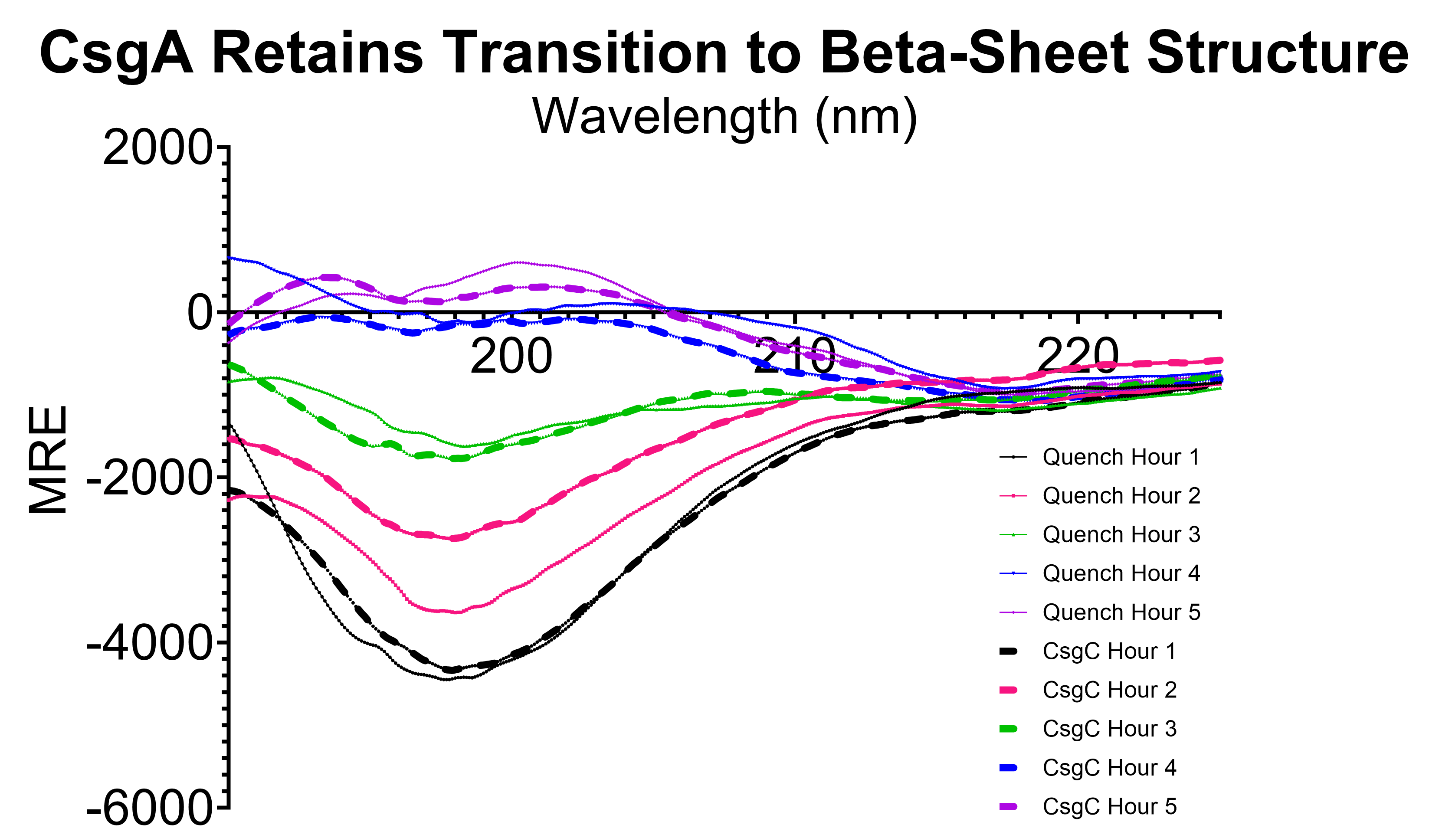

**Fig. S8. Circular Dichroism of CsgA eluted from quenched or CsgC-linked resin.** Circular dichroism of 15 µM CsgA eluted from quenched NHS-Resin (called Quench Hour X, solid lines) or CsgC-linked resin (called CsgC Hour X, dashed lines) X hours following purification. Measurements were taken in triplicate and averaged. Measurements were also taken over two independent assays to account for sampling order.

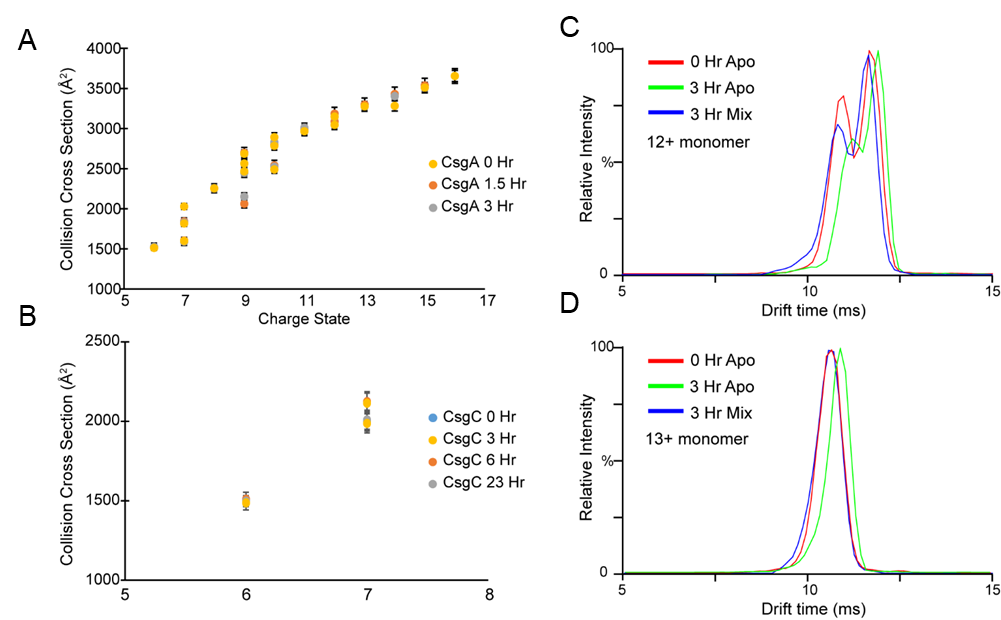

Fig. S9. Additional IM-MS Data. CCS values derived from arrival time distribution of A) CsgA monomers and B) CsgC monomers. C and D) Arrival time distribution of the 12+ and 13+ monomer of CsgA in three experimental conditions: Apo CsgA in solution at 0 hr (red trace), apo CsgA in solution after 3 hr (green trace), and CsgA in solution with CsgC after 3 hr (blue trace). Traces are overlapped to show the changes in ATD after 3 hr of incubation.

**Supporting Tables**

Supporting Table 1. Strains used in this study.

| Strains | Relevant Genotype | References |
| --- | --- | --- |
| NEB3016 | MiniF *lacI^q^*(Cam^R^) */ fhuA2 lacZ::T7 gene1 [lon] ompT gal sulA11 R(mcr-73::miniTn10--*Tet^S^*)2 [dcm] R(zgb-210::Tn10--*Tet^S^*) endA1 Δ(mcrC-mrr)114::IS10* | New England Biolabs |
| MC1061 | *F– araD139 Δ(ara-leu)7696 galE15 galK16 Δ(lac)X74 rpsL (StrR) hsdR2 (rK– mK+) mcrA mcrB1* | (55) |
| CsgA | NEB 3016 ΔslyD + pCsgA | (Evans, 2015) |
| CsgC EC | NEB 3016 ΔslyD + pCsgC EC | (Evans, 2015) |
| RAL1 | NEB 3016 ΔslyD + pCsgC CY | This study |
| RAL2 | NEB 3016 ΔslyD + pCsgC CD | This study |
| RAL3 | NEB 3016 ΔslyD + pCsgC HA | This study |

Supporting Table 2. Plasmids used in this study.

| Plasmids | Relevant Characteristics | References |
| --- | --- | --- |
| pET11d | IPTG inducible expression vector | New England Biolabs |
| pET28a | IPTG inducible expression vector | New England Biolabs |
| pCsgA | pET11d-CsgA -sec 6xHis | (Zhou, 2012) |
| pCsgC EC | pET28a-CsgC-sec C-term 6xHis, kan^r^ | (Evans, 2015) |
| pCsgC CY | pET28a-*Citrobacter youngae* CsgC-sec C-term 6xHis | This study |
| pCsgC CD | pET28a-*Cedecia davisae* CsgC-sec C-term 6xHis | This study |
| pCsgC HA | pET28a-*Hafnia* *alvei* CsgC-sec C-term 6xHis | This study |

Supporting Table 3. Primers used in this study.

| Primers | Primer Sequence (5’→ 3’) | Constructs |
| --- | --- | --- |
| GA CsgC CY Vector_FOR | ATTTGATTTGCCAGCGCCATGGTATATCTCCTTCTTAAAGTTAAACAAAATTATTTCTAG | pCsgC CY |
| GA CsgC CY Vector_REV | CTGACTCAACCCATAGCCCACACCACCACCACCACCACT | pCsgC CY |
| GA CsgC CY Frag_FOR | CAGTGGTGGTGGTGGTGGTGTGGGCTATGGGTTGAGTCAGGC | pCsgC CY |
| GA CsgC CY Frag_REV | TAAGAAGGAGATATACCATGGCGCTGGCAAATCAAATAACATTCAA | pCsgC CY |
| GA CsgC CD Vector_FOR | agcactaaggtatgcatGGTATATCTCCTTCTTAAAGTTAAACAAAATTATTTCTAGAGG | pCsgC CD |
| GA CsgC CD Vector_REV | gggtcggggcaggaaagccgCACCACCACCACCACCACT | pCsgC CD |
| GA CsgC CD Frag_FOR | CTTTAAGAAGGAGATATACCatgcataccttagtgctgcttgc | pCsgC CD |
| GA CsgC CD Frag_REV | CAGTGGTGGTGGTGGTGGTGcggctttcctgccccgac | pCsgC CD |
| GA CsgC HA Vector_FOR | CAGAGCTGATTAGATAAAACCATGGTATATCTCCTTCTTAAAGTTAAACAAAATTATTTCTAGAG | pCsgC HA |
| GA CsgC HA Vector_REV | TGATACTACCCGATAGGGTTCACCACCACCACCACCACT | pCsgC HA |
| GA CsgC HA Frag_FOR | CAGTGGTGGTGGTGGTGGTGAACCCTATCGGGTAGTATCACTTGT | pCsgC HA |
| GA CsgC HA Frag_REV | TAAGAAGGAGATATACCATGGTTTTATCTAATCAGCTCTGGTTCGATACCAC | pCsgC HA |
